# Optimized optogenetic anti-CRISPR for endogenous gene regulation in *Drosophila*

**DOI:** 10.1101/2025.05.02.651826

**Authors:** Clarissien Ramongolalaina, José C. Pastor-Pareja, Emily Zhang, Yichang Jia

## Abstract

Optogenetic tools – whose engineering requires a structural understanding of the target proteins – are attractive approaches for achieving endogenous gene regulation under minimally invasive conditions. To build an optogenetic system for controlling endogenous gene expression, we first identified Anti-CRISPR (Acr) proteins that can inhibit CRISPRa-mediated transcriptional activation in *Drosophila*. Next, we inserted optogenetic protein LOV2 in these Acrs, tested for their ability to optogenetically modulate endogenous gene upregulation (EnGup) through the CRISPRa-based flySAM system in *Drosophila*, and found that the photoswitchability of these prototypes was weak. Hence, we engineered an Acr-LOV2 fusion module with refined length of intrinsically disordered and ordered regions (IDR and IOR) and optimized LOV2. This optimized variant, whose application yielded new findings *in vivo*, was significantly more sensitive for EnGup under blue light than the prototypes. In short, not only does this work introduce the application of Acr proteins in an animal model, but it also provides insights for *in vivo* characterization of the IDR and the IOR of these small-sized proteins. Together, these findings establish a robust optogenetic toolbox for precise, light-sensitive endogenous gene regulation in *Drosophila*.

**Graphical abstract:** 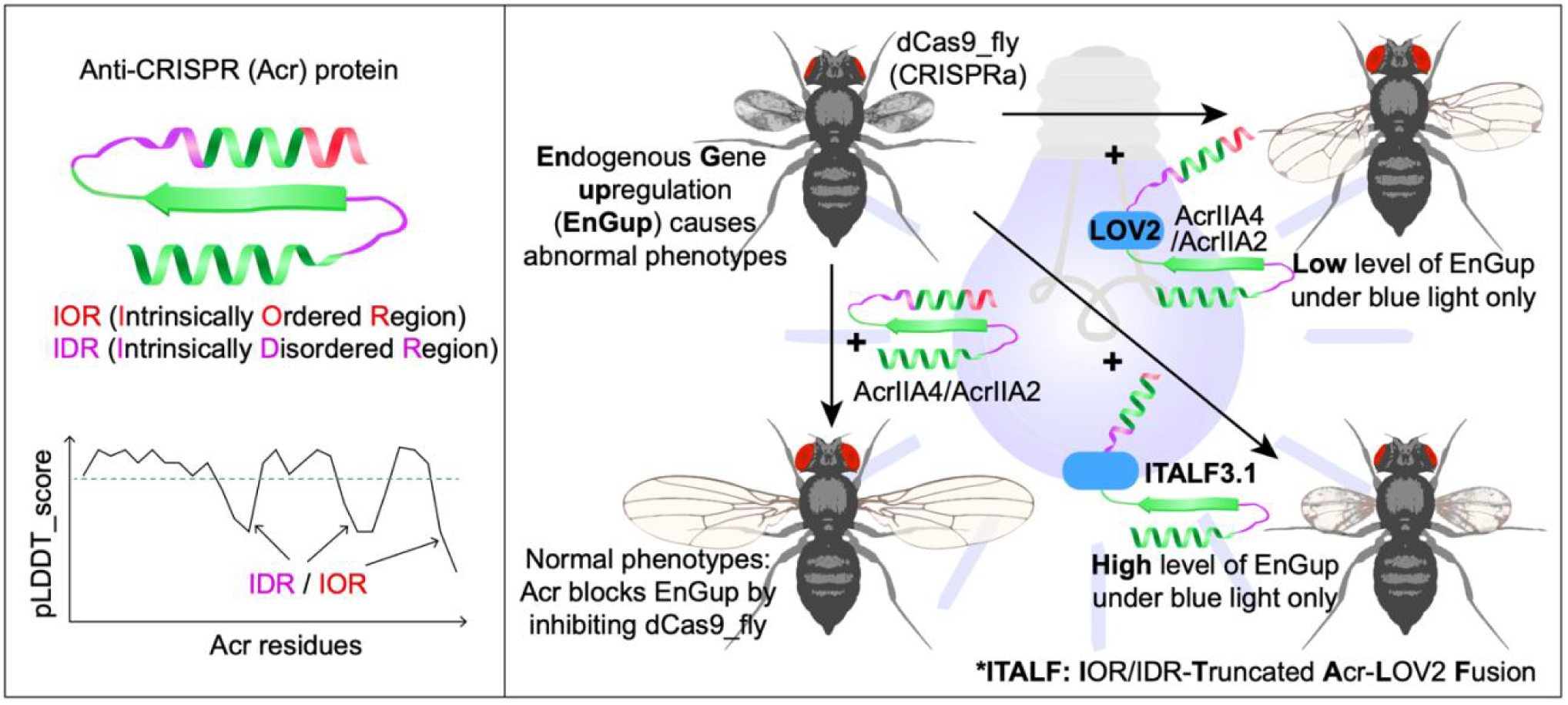

## INTRODUCTION

Optogenetics, the integration of optics and genetics, can be defined as an approach to control the function of an ‘opsin’ protein using light. Originally a cornerstone of neuroscience, it has been also harnessed by the biotechnology field to spatiotemporally modulate gene expression, protein interactions and countless biological reactions not exclusively for research purposes, but for therapeutic applications as well (1). Its implementation to regulate endogenous gene expression *in vivo*, is promising; this owes to light features such as sharpness, spatial resolution (2) and spectral multiplexing (3). In a model organisms such as *Drosophila*, existing systems enable efficient optogenetic control of genes or proteins are expressed from transgenes (4–6). In contrast, endogenous gene regulation in *Drosophila* is currently achievable via CRISPRa (7) – hereafter named dCas9_fly – and CRISPRi (8), but these systems lack inherent light-switchability.

In engineering light-switchable tools, understanding the detailed structure of the protein of interest, more distinctively its intrinsically disordered regions (IDRs), is critical. IDRs are ideal switch sites for insertion of an opsin (9), although IDR-mediated interactions often differ between *in vitro* and *in vivo* systems, as demonstrated in yeast and mouse models (10–15). IDR – a peptide that has multiple conformations, without 3D-structure (16) – is generally a crucial part of any given proteome, and is associated with many functions, including cell signaling and protein interactions (17), and the formation of membraneless organelles (18). Accordingly, several approaches have been developed for the prediction of IDRs; amongst those proven to be accurate is the alphaFold2-defined pLDDT scores of protein models (19–21). Intriguingly, the theoretical computation of pLDDT scores leads to the identification of 3D-folded IDRs (21) that would not be otherwise classified as IDRs and have never been experimentally characterized. New web servers such as MobiDB (22) have been continuously emerging for prediction and characterization of IDRs; however, they do not provide structural details of small size proteins such as anti-CRISPR (Acr) proteins.

Acr bacteriophage proteins act as natural antagonists of CRISPR-Cas systems. Amongst the 122 non-homologous Acr proteins discovered to date, and classified into four types, the type II families can inhibit DNA cleavage by CRISPR-Cas9 (23). In doing so, Acr proteins, controlled with opsins, can serve as proper tools to optogenetically modulate CRISPR-Cas9 activities. The LOV2 photoreceptor from *Avena sativa* (oat) is the most used opsin in research. In the presence of the chromophore flavin mononucleotide (FMN), a cofactor that exists in almost all species, it can be optimally and reversibly activated by 450 nm blue light (24). In the last few years, type II Acr proteins inserted with LOV2 have shown CRISPR-Cas9 light-switchability *in vitro* (25, 26) and in *E.coli* (27).

Despite the promise of optogenetic CRISPR for endogenous gene regulation *in vivo*, several key challenges remain: Can Acr protein be functional in *Drosophila*? If so, how to engineer a photoswitchable Acr without disrupting its functionality? How would the functional significance of Acr IDRs, the binding affinity of which *in vitro* and *in vivo* may differ, come into play? And how can this system be optimized to achieve light-sensitive phenotypic effects? Addressing these points herein, we identified the Acr proteins that can inhibit dCas9_fly-mediated EnGup in *Drosophila*, analyzed structures of these Acrs to determine their LOV2 insertion sites and generated flies expressing Acr-LOV2 fusion prototypes, which we next optimized mainly by refining the length structures that we defined as intrinsically disordered and ordered regions (IDR and IOR) in Acrs. Finally, we demonstrated the application of our optimized system in autophagy investigation and phenotypic modeling through manipulation of endogenous gene expression.

## RESULTS

### 1. AcrIIA4a and A4b efficiently deactivate dCas9_fly-mediated EnGup in *Drosophila*

First, we selected five type II Acr family members that can inhibit CRISPR-Cas9: AcrIIA2 along with AcrIIA4a, AcrIIA4b and AcrIIA6 – inhibitors of SpyCas9 (28, 29) – from *Listeria monocytogenes*, *L. farberi* and *Caudoviricetes sp*., respectively; and AcrIIC3 – an antagonist of NmCas9 (25) – from *Neisseria meningitidis*, as a negative control (Figure S1A). We generated UAS constructs to express each Acr protein under control of the Gal4-UAS binary expression system and introduced them transgenically into flies at the *attP40* site on chromosome II. To assess potential toxicity of the expression of individual Acr proteins *in vivo*, we crossed each generated Acr line with generally expressed Gal4 driver Tub-Gal4, and tissue-specific drivers Mef2-Gal4 (muscle) and BM-40-SPARC-Gal4 (adipose tissue). Although the expression of AcrIIA6 by these three Gal4 drivers caused a significantly shorter lifespan compared to that of controls, the expression of AcrIIA2, AcrIIA4a, AcrIIA4b, or AcrIIC3 produced normal survival rates (Figure S1B-1D). In addition, no visible morphological or developmental defects were observed in these animals (Figure S1E), which allowed us to continue our studies *in vivo* by expressing these Acr proteins.

Next, we evaluated which Acr proteins could effectively inhibit dCas9 from binding DNA, and thereby prevent dCas9_fly-mediated EnGup *in vivo* (Figure 1A). To that end, we crossed each Acr line flies with dCas9_fly targeted different genes: *dpp*, *vn*, *upd2*, *foxo*, and *wnt4* (Figure 1B). The transcriptional activation of those endogenous target genes by dCas9_fly was detected as previously described (7): mRNA expressions from dCas9_fly were higher than from negative control (Figure 1C). Of note, although EnGups *in vivo* of the five genes were relatively similar at the mRNA (Fig 1C), those of *dpp* and *vn* produced very consistent and strong phenotypes in eyes and particularly in wings (Figure 1D-1G). More importantly, the inhibition of dCas9_fly-mediated EnGup *in vivo* was evidenced at the mRNA level, since upregulation of the five endogenous target genes was inhibited by expression of AcrIIA2, AcrIIA4a, or AcrIIA4b, but not AcrIIC3 (Figure 1C). Interestingly, supported by phenotypic effects, AcrIIA4a and A4b consistently demonstrated stronger inhibition than AcrIIA2: the eye and wing defects induced by dCas9_fly activation were largely reversed by coexpression of AcrIIA4a or A4b, but partially by AcrIIA2 (Figure 1D-1G and Figure S2A-S2E). Despite its lower efficiency than AcrIIA4a and A4b, AcrIIA2 could be more efficiently inhibit Cas9, specially at a low temperature condition, in *Drosophila* than in other species (Figure S2F-S2G); AcrIIA2 barely inhibits Cas9 in mammalian cells and at 37°C, and reduced only up to 25% of Cas9 activities in *E.coli* as previously demonstrated (30, 31). Together, these results demonstrate that AcrIIA4a and A4b effectively suppress dCas9_fly-mediated EnGup, and AcrIIA2 could potentially do so *in vivo Drosophila*.

**Figure 1.**
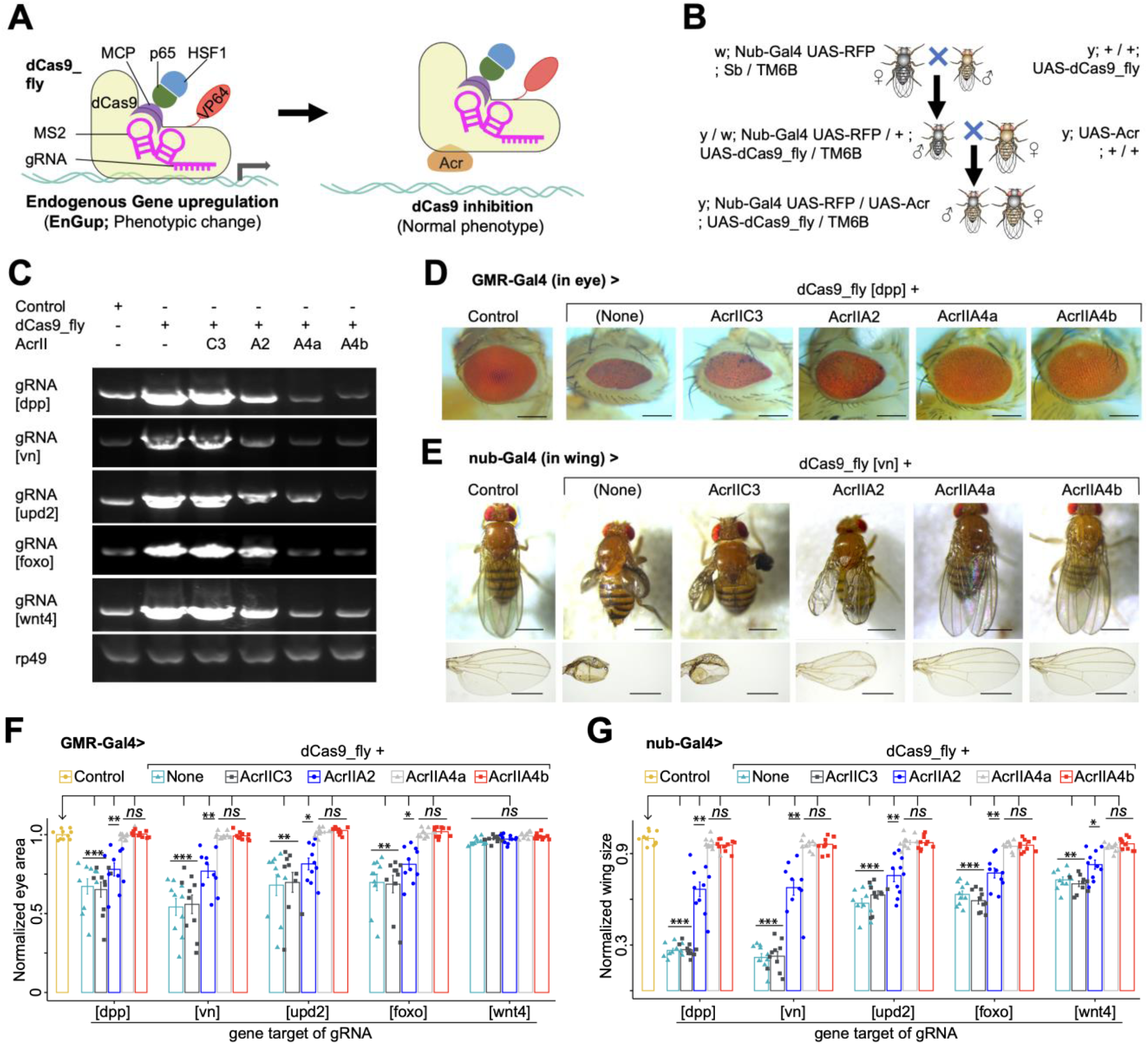
Identification of Acr that effectively inhibits dCas9_flySAM. (**A**) Cartoon representation of dCas9_flySAM for target gene expression in flies. The system is composed of dCas9 fused to transcriptional activation domain p65-HSF1, via MS2-MCP, and to a chimeric activator VP64, directly. Acr protein is hypothetically able to block dCas9_flySAM activity. (**B**) Crossing scheme for coexpressing dCas9_flySAM and Acr with a non-lethal driver, such as Nub-Gal4 in wings. (**C**) RT-PCR results of fly wings that expressed dCas9_flySAM-targeted a set of genes (*dpp*, *vn*, *upd2*, *foxo* and *wnt4*) along with each Acr protein. *rp49*, a loading control. (**D**-**G**) Representative images showing that AcrIIA4a&b are effectively able to rescue the phenotypes generated by *in vivo* CRISPRa via inhibition of dCas9 (**D** and **E**) and corresponding data summaries were shown (**F** and **G**). Data summaries for eye surfaces (**F**) and wing sizes (**G**) of flies coexpressing dCas9_flySAM-targeted genes (*dpp*, *vn*, *upd2*, *foxo* and *wnt4*) along with each Acr protein. In **F** and **G**, every single value is normalized with the mean of normal control. Values are represented as means ± s.d.; n = 10 female flies. *P* values were calculated by ANOVA-test; *ns*, *, ** and *** are respectively not significant, <0.01, <1e-3 and <1e-5 compared to the normalized control phenotypes. In **D** and **E**, scale bars are 500 μm and 250 μm, respectively.

### 2. The residues adjacent to IDR are predicted to be the optimum sites for opsin LOV2 insertion

To engineer a light-switchable system, we identified optimal insertion sites for the *Avena sativa* LOV2 photoreceptor in AcrIIA4 and AcrIIA2 (the potent one) by analyzing the predicted structures of these Acr proteins (Figure 2A). Structural analysis using AlphaFold3 revealed that although the two AcrIIA4 primary structures are 40% dissimilar, they possess nearly identical tertiary structures and hydrophobicity patterns (Figure 2B, Figure S3A-S3C). However, the charge distribution of their residues (Figure S3D) and their electrostatic potentials on the surface differ slightly (Figure 2C). More explicitly, although their surfaces are acidic, a common feature of AcrIIA4 family (32), F⍺ helix of AcrIIA4a is highly negative compared to that of AcrIIA4b. As for AcrIIA2, its structure have little resemblance with those of AcrIIA4 family (Figure 2B and S3A), because the molecular mechanisms by which these Acrs block DNA binding of Cas9 are different (30, 33).

**Figure 2.**
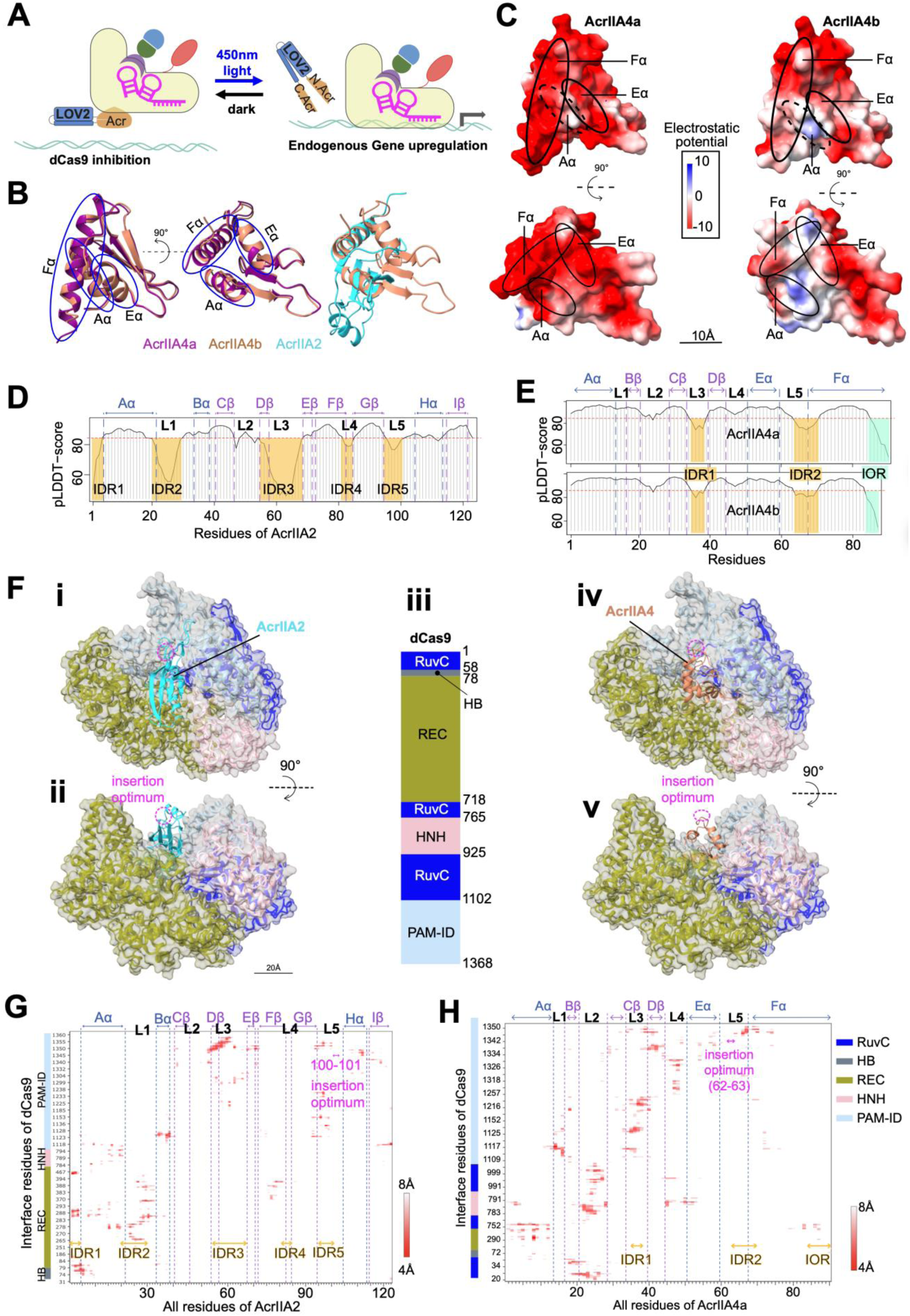
Structural analyses of Acr proteins. (**A**) Schematic of the system that we intended to build in Drosophila to enable gene overexpression only under the light illumination. (**B**) Side views of ribbon representation of AcrIIA4a, AcrIIA4b (homology PDB ID 5VW1) and AcrIIA2 (homology PDB ID 6MCB), colored in warm purple, light caramel and bright cyan, respectively, captioning the structural similarities of the two AcrIIA4(s). The first view (left) is rotated to 90° angle in the second (middle) along the vertical axis (dashed line). The alpha-chains of AcrIIA4(s) (Aα, Eα, Fα) are highlighted. (**C**) Top (left) and side (right) views of electrostatic potentials of the molecular surfaces of AcrIIA4a (upper) and AcrIIA4b (bottom). (**D**-**E**) The predicted local distance difference test (pLDDT) scores of AcrIIA2 (D) and AcrIIA4a&b (E). Their IDRs and IORs below the threshold (pLDDT_score= 85, in red dashed-line) are labeled and spotlighted, respectively, in semitransparent orange and turquoise. (**F-G**) Predicted interaction of AcrIIA2 and AcrIIA4b with dCas9. (**F**) Top (**i**&**iv**) and front (**ii**&**v**) views of alfaFold model dCas9 in complexity with AcrIIA2 (left; homology PDB ID 6IF0) and AcrIIA4 (right; homology PDB ID 5VW1). (**iii**) Schematic diagram of domain organization of SpyCas9. The RNA recognizing lobe (REC), two nuclease domains HNH and RuvC, and PAM-interacting domain (PAM-ID) of the dCas9 domains are colored according to the scheme in (iii). (**G-H**) Contact maps of predicted the interface residues of dCas9 with AcrIIA2 (**G**) and AcrIIA4a (**H**) within 8Å distance of interactions. The α-chains, β-sheets and loops (L1-5) of AcrIIA4a are labeled. Likewise, the IDRs and IOR are highlighted in orange. In **F, G** and **H**, the optimum insertion sites are spotlighted in magenta.

As IDRs play key roles in protein engineering (34, 35), we computed the pLDDT scores – outlined by alphaFold3, formerly by alphaFold2 (36) – of AcrIIA2 along with AcrIIA4a and A4b. Due to the small size of Acr proteins, we defined IDRs as four amino acid sequences or above without 3D structures, having the pLDDT score < 85, similar criteria was reported previously (16). Notably, with this computation of pLDDT scores, there are peptide sequences – at lease 4 aa in length, with pLDDT < 85 and 3D-structures – that would not be otherwise classified as IDRs (16) and formerly proposed as 3D-folded IDRs (21); we named Intrinsically Ordered Regions (IORs) these peptide sequences. In another words, an IOR is four amino acid sequences or above with the pLDDT score < 85, and natively possesses 3D-structure. With these defined IDRs and IORs, our analysis displays that AcrIIA2 is IDR-enriched (∼ 30% of full sequence), 25%, 40% and 20% among IDRs are around the loops L1 (21-29 aa), L3 (55-68 aa) and L5 (95-100 aa), respectively (Fig 2D). Alternatively, AcrIIA4a and AcrIIA4b have short IDRs that extend in loops L3 (35-38 aa) and L5 (64-70 aa), alongside an IOR at their C-terminus, which is seven residues (84-90 aa) for A4a and four (84-87 aa) for A4b (Figure 2E). The IDRs and IORs of these three Acr proteins contain a proportion of acidic residues. This less-hydrophobic feature of IDRs is consistent with the pLDDT-score-defined IDRs in big size proteins (19–21), even though Acr proteins are small in size (< 125 aa).

To determine optimal LOV2 insertion sites – residues in a loop of Acr protein that are not in close contact with any dCas9 residue (distance from the dCas9 interfaces > 8Å) – we generated AlphaFold-predicted Acr-dCas9 complex models (Figure 2F). To this end, we performed intercontact map analyses of these alphaFold models, in which AcrIIA2 or AcrIIA4 blocks the DNA binding pocket of dCas9, who possesses several domains (Figure 2F). These analyses reveal that, as previously reported (15), AcrIIA2 interfaces with mainly the PAM-identification and RNA-recognizing lobe of dCas9 (Figure 2G), whereas AcrIIA4a and A4b do so with every domain of the latter (Figure 2H and S3E) – explaining their robust inhibition of dCas9_fly (Figure 1). Interestingly, IDRs of AcrIIA4a and A4b more strongly interact with dCas9 than their IORs, indicating that IDRs are functionally important for the interactions and not suitable for opsin LOV2 insertion (Figure 2H and S3E). Inferring from these Acr-dCas9 predicted interface regions, the residues 100-101 (Loop L5, adjacent to IDR5, distance from dCas9 interface > 8Å) of AcrIIA2 and 62-63 (loop L5, adjacent to IDR2, distance from dCas9 interface > 8Å) of AcrIIA4a along with A4b are more likely the optimum sites for LOV2 insertion (Figure 2G, 2H and S3E). On that account, our predicted insertion sites of LOV2 did not support the previous findings, proving that IDRs are the optimum sites of insertion of an opsin (9, 26). Taken together, for these *in silico* analyses, not only did we predict optimal insertion sites of LOV2 in AcrIIA2 and AcrIIA4, which are rather adjacent to IDRs, but we also proposed the existence of IOR in these small proteins.

### 3. A4b.LOV_63 a prototype photo-switchable component functional *in vivo*

Based on our predicted optimum insertion of LOV2, we built UAS plasmids expressing the prototypes of Acr-LOV2 fusion proteins and introduced them transgenically into flies in order to validate photo-switchable endogenous gene regulation *in vivo*. More notably, LOV2 – whose gap between C- and N-termini is ∼10Å in the dark, and C-terminal Jα helix unfolds upon photoactivation (9, 37) (Figure S4A) – was inserted into four sites of A4a and A4b: N25, Q47 (of A4a) and E47 (of A4b), G62 (of A4b) and W63 (of A4a), and E66 (Figure 3A, 3B, S4B, and S4C). The N25 (in Loop2) and Q47/E47 (in Loop4) close to the AcrIIA4-dCas9 interface act as negative controls. Previously reported optimum insertion site E66 was included as a positive control (26) (Figure 3A, 3B, S4B, and S4C). In this case, the optimal site amongst these positions was picked out by the ability of Acr-LOV2, when coexpressed with dCas9_fly, to procure a maximum dark-light phenotypic difference: the intact Acr protein that completely blocks dCas9-mediated phenotypic changes is split by LOV2 under blue light, permitting endogenous gene activation (Figure 3C). Before the dissections of the wings from adult flies for phenotypic analyses, all breeding processes were carried out in the dark; otherwise, pupae or larvae were illuminated under a blue light pulse (Figure 3D).

**Figure 3.**
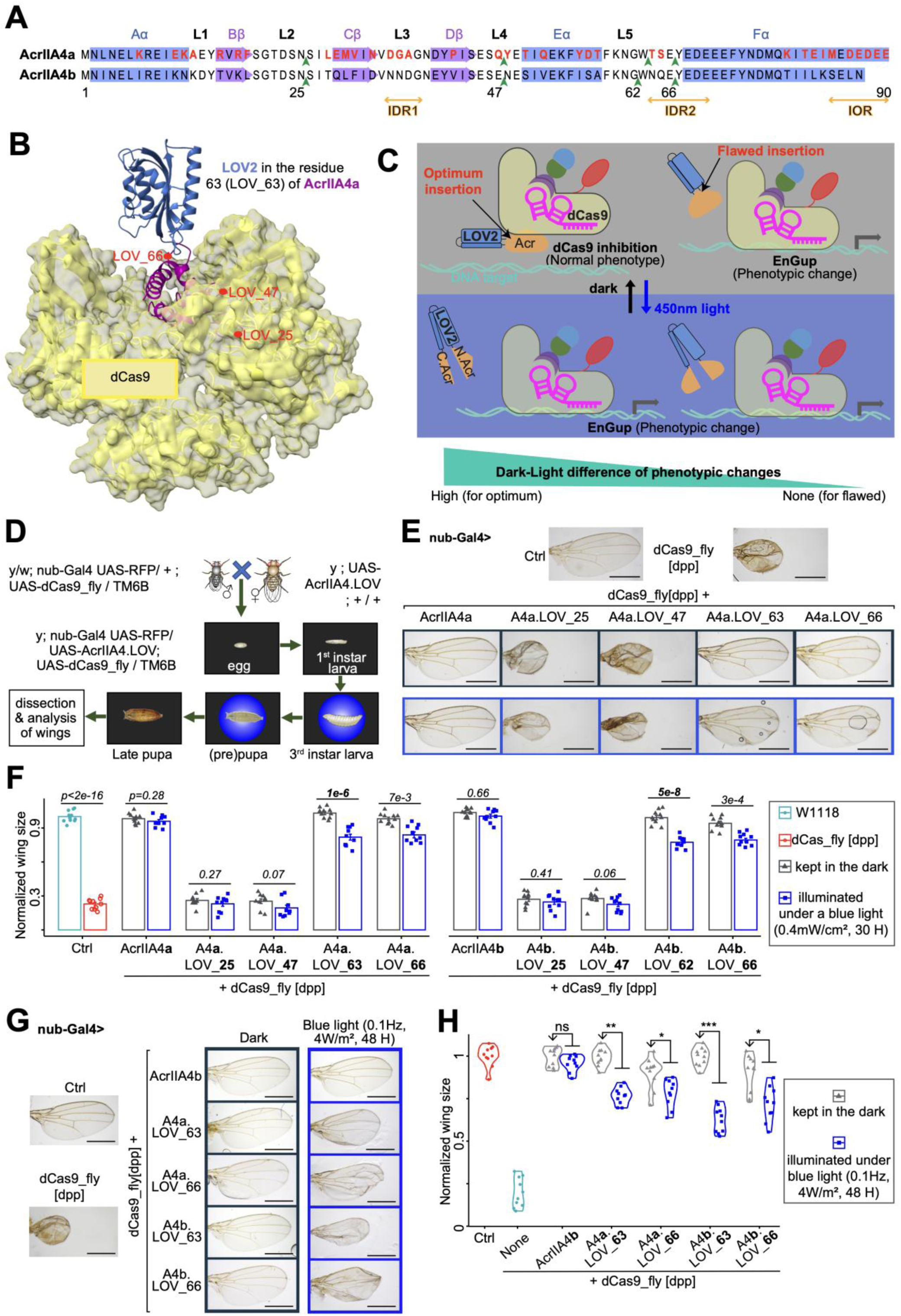
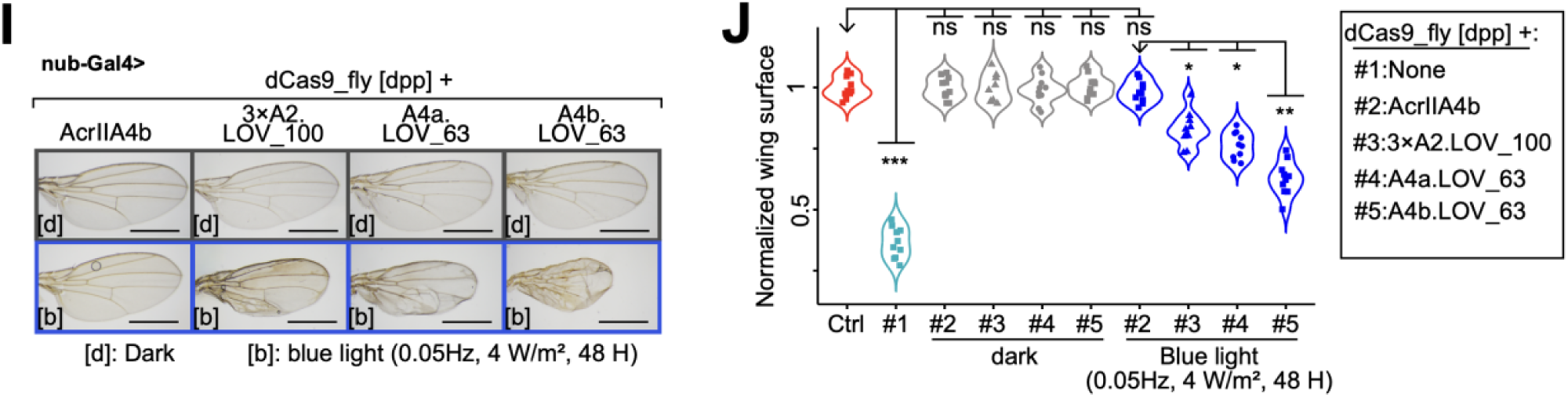
**Engineered Acr proteins and their light-switchable to modulate gene overexpression *in vivo***. (**A**) Representations of secondary structure of AcrIIA4a and aligned with that of AcrIIA4b, and their selected LOV2 insertion sites (in green arrowhead). The amino acid sequences of alpha-chains, beta-sheets and loops are colored in blue, medium orchid and black, respectively. The IDRs and IOR are marked in orange. The red characters are the residues differentiate AcrIIA4a from A4b. (**B**) Predicted structural model of the system for the LOV2 (dark blue) inserted in W63 of AcrIIA4a. The other LOV2 insertion sites are also highlighted in red. The surface of dCas9 is colored in semi-transparent yellow. (**C**) Rationale behind the optimum site of LOV2 insertion for inhibition of dCas9_fly. When LOV2 (dodger blue) is inserted in the optimum position of Acr protein (orange), the latter is able to inhibit dCas9 in dark but is split by the active LOV2 under blue light allowing dCas9_fly-mediated EnGup. (**D**) Experimental workflow. The crossing and their progenies were kept in the dark, temporarily illuminated with a blue LED-light at the 3rd instar larval, prepupal or pupal stage, and put back in the dark before the dissection of the wings of adults. (**E**) Illustrative images of wings from flies coexpressing dCas9_fly-mediated *dpp* EnGup and LOV2-inserted AcrIIA4 at different positions. The blue-framed images were from the larvae exposed to a blue LED-light. (**F**) Data summary of normalized wing sizes described in **E**. The numbers on the top of each pair of bars are the *p* values of ANOVA tests of the dark-light difference with n = 10 female flies. (**G**) Representative images showing the light-responsiveness of LOV_63 of A4a and A4b compared to the positive control LOV_66. (**H**) Data summary of normalized wing sizes described in **G**. It displays that LOV_63 of A4a and A4b is more light-responsive that LOV_66 for *in vivo* drosophila. (**I-J**) Representative images (**I**) and summarized data (**J**) illustrate the light sensitivity of AcrIIA4b compared with AcrIIA4a and triple dose AcrIIA2. In **E**, **G** and **I**, scale bars are 500 μm. The dark gray- and blue-framed images are the wings from entirely kept in the dark and temporarily illuminated with a blue LED-light, respectively. In **H** and **J**, *P* values were calculated by ANOVA-test; n = 10 female flies; *ns*, *, ** and *** are respectively not significant, <0.01, <1e-3 and <1e-5 between the normalized phenotypes of the arrowhead-pointed group and those of other groups.

After checking the wing phenotypes of adult flies whose larvae were kept in dark or exposed to blue light, we expectedly found that LOV_25 and LOV_47 of A4a and A4b, whether in dark or light conditions, exhibited phenotypes similar to dCas9_fly alone (Figure 3E and 3F), suggesting that these two LOV2 insertions in A4a and A4b disrupted A4 protein inhibitory function *in vivo*. The LOV2 insertions in W63 of A4a and in G62 of A4b, together with that in E66 of both AcrIIA4, were light-responsive, because wings were significantly smaller when larvae were exposed to a blue light (0.1Hz, 0.4mW/cm^2^) than when kept in the dark (Figure 3E and 3F). Assessing the two adjacent positions (G62 and W63) in A4b, LOV_63 performed significantly better than LOV_62, as the G62 site is one residue apart from the IDR2 (Figure S5A and S5B). Different from LOV_25 and LOV_47 insertions disrupting inhibitory function of A4 (Figure 3E and 3F), LOV_62 and LOV_63 insertions in A4b had little effect on wing size in dark conditions compared to that of A4b alone, indicating that both insertions of LOV2 keep A4b protein functionally intact (Figure S5A and S5B). The in-depth examination of the insertions in W63 and E66 of A4a and A4b exhibited that A4b.LOV_66 defectively inhibited dCas9 in the dark, leading to less consistent wing sizes, and A4b.LOV_63 produced a maximum dark-light phenotypic difference (Figure 3G and 3H).

Based on our prediction (Figure 2G), we also generated fly lines expressing A2.LOV_100 as a potentially light-switchable fusion and included A2.LOV_45 (T45 in Cß, Figure S3B) as a negative control (Figure S6A). Using genetic manipulation, we doubled the expression of AcrIIA2 (2×A2) (Figure S6B). As expected, 2×A2.LOV_45 disrupted inhibitory effect of AcrIIA2; however, 2×A2.LOV_100 significantly decreased the wing size when fruit flies were exposed to blue light (0.1Hz, 4W/m^2^), compared to those in dark condition (Figure S6C and S6D). To further dose A2.LOV_100, we generated a fly line expressing tricistronic A2.LOV_100 (3×) that was able to more effectively inhibit dCas9_fly than 2×A2.LOV_100 in dark condition (Figure S6E-S6G). Even so, the dark-light differences of 3× and 2×A2.LOV_100 were similar, meaning that the photoswitchability of Acr-LOV2 fusion did not depend on the dosage of AcrIIA2 (Figure S6G). Subsequent comparison of 3×A2.LOV_100 with LOV_63 of the two A4 variants displayed that these three variants were able to effectively inhibit dCas9_fly in dark, and were light-responsive (Figure 3I and 3J). A closer look, however, showed that A4b.LOV_63 amongst these three variants was more photoswitchable (Figure 3I and 3J). In short, although the photoswitchability needed to be further improved, A4b.LOV_63 was a prototype for photo-switchable EnGup with dCas9_fly *in vivo*.

### 4. Refining the length of IDR and IOR of A4b.LOV_63 increases its photoswitchability *in vivo*

Since AcrIIA4a has longer IOR than A4b and the insertion of LOV2 in AcrIIA2 (L100) is in the IDR5, we reasoned that residues in IDR and IOR of A4 proteins would provide an opportunity to achieve the optimization of our prototype photo-switchable A4b.LOV_63 *in vivo*. Realizing that IDR of AcrIIA5 was previously reported to be crucial for Cas9 inhibition (38), we created triple-residue-deletion variants (Figure 4A, S7A and S7B): Δ3, Δ4 as a negative control, Δ5, Δ6, double truncated Δ7’ and partially double truncated Δ8 (∂T64/E89/E90 for A4a and ∂N64/L86/N87 for A4b). These variants aimed to identify truncatable residues in IDRs and IOR of A4.

**Figure 4.**
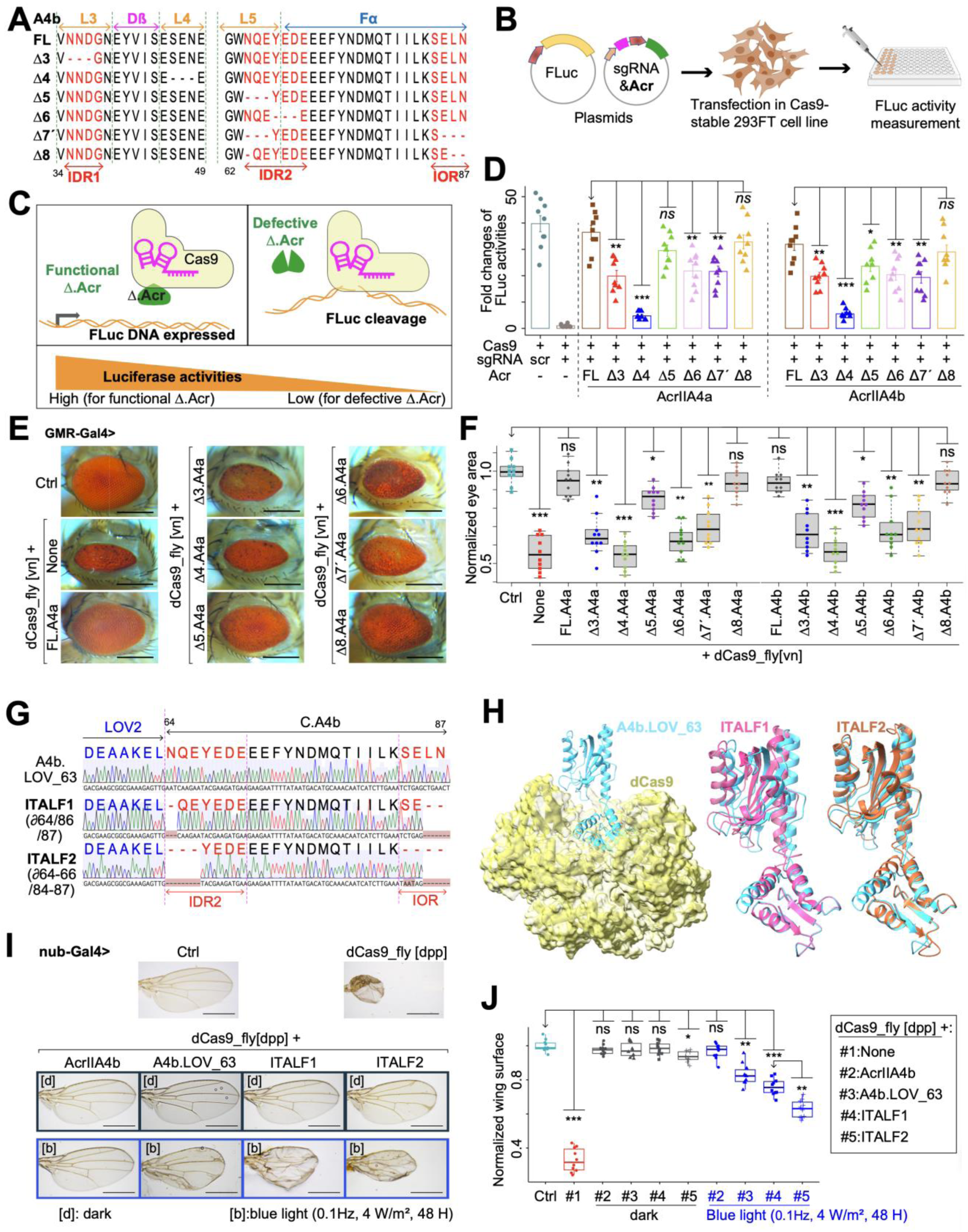
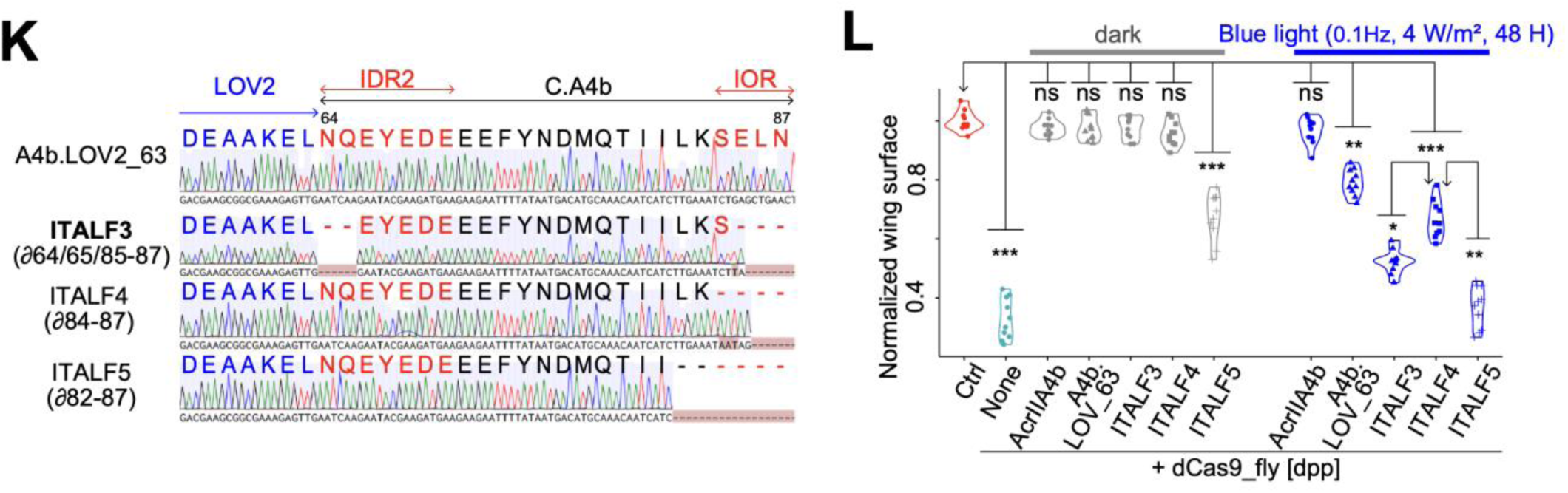
Optimization of A4b.LOV_63 photoswitchability *in vivo* by truncation of IDR2 and IOR. (**A**) Partial amino acid sequences of truncated AcrIIA4b variants aligned with the full-length one (FL). Δ7’ indicates a double truncation, and Δ8 is deletion of tree residues in AcrIIA4b (∂64/86/87). The secondary structures are outlined. (**B**) Experimental design for testing the inhibitory efficiency of truncated Acrs. Cells stably expressing Cas9 were cotransfected with each truncated Acr, sgRNA targeting Luciferase, and a reporter Firefly Luciferase. (**C**) Rationale behind the identification of truncatable residues of Acr: high Luciferase activity indicates the truncated residues are not necessary for the Cas9-inhibiting functionality of Acr variant, and vice-versa. (**D**) Plot showing the results of Firefly luciferase activities targeted by Cas9 according to the inhibition efficiency of co-transfected Acr. The AcrIIA4a&b are truncated corresponding to **A** and Figure S7A-B; ‘scr’ means scrambled sgRNA. Bars represent mean values, error bars the standard deviation and dots individual data points from n =9. (**E**) Representative images of compound eyes from *Drosophila* coexpressing dCas9_fly and FL or truncated AcrIIA4a&b proteins mentioned in **A** and Figure S7A-B. (**F**) Quantitative data of eye surface area from dCas9_fly drosophila expressing truncated AcrIIA4A&b. (**G**) Partial sequence graphs of IOR/IDR-truncated Acr-LOV2 complexes ITALF1 and ITALF2 aligned with A4b.LOV_63, featuring their truncated residues ∂64/86/87 and ∂64-66/84-87, respectively. (**H**) Generated alphaFold of ITALF1 (magenta) and ITALF2 (coral), aligned with A4b.LOV_63 (cyan), which interacts with dCas9 (yellow). (**I-J**) Representative images (**I**) and statistic data (**J**) of wing phenotypes from dCas9_fly flies expressing Acr-LOV2 complexes in **G** and **H**, and kept in the dark or exposed to blue-light pulse at the pupal stage. (**K**) Partial sequence graphs and corresponding peptide sequences of ITALF3, ITALF4 and ITALF5, whose residues 64-65/85-87, 84-87 and 82-87, respectively, are truncated, and aligned with A4b.LOV_63. (**L**) Statistic data set of normalized wing sizes from dCas9_fly drosophila expressing Acr-LOV2 complexes in **K**, displaying the high and marginal photoswitchabilities of ITALF3 and ITALF4, respectively, and disrupted functionality of ITALF5. The scale bars in **E** and **I** are respectively 250 μm and 500 μm. In **A**, **G** and **K**, the sequence graphs were aligned with Benchling platform; the dashes or codons in blossom color are deletion or mismatches. The residues in blue and red are LOV2 and IDR2 plus IOR of C.AcrIIA4b. The dash (-) represents a single residue deletion. In **D**, **F**, **J** and **L**, the *p* values * < 0.01, ** <1e-3, *** < 1e-5 were calculated by one-way analysis of variance (ANOVA), with n = 10 female flies (for **F**, **J** and **L**), of the differences between the normalized phenotypes of the arrowhead-pointed group and those of other groups. In all panels, ∂ (lower case ‘Delta’) and Δ (upper case ‘Delta’) refer to single and more than one residue deletions, respectively.

To assess their functionality, we cotransfected these truncated mutants, at first, with Cas9 plus sgRNA along with *Firefly Luciferase*, the sgRNA-targeted gene, in 293FT cells (Figure 4B). In this case, Luciferase activity served as a reporter for *Luciferase* DNA cleavage by Cas9 when at low activity level, and Cas9 inhibition by an A4 mutant when at high level (Figure 4C). Luciferase assays revealed that all deletion mutants were relatively inefficient to block Cas9, except two variants: Δ4 was completely dysfunctional, and Δ8 effectively inhibited Cas9, latter of which was comparable to full-length (FL) A4a or A4b group (Figure 4D). These results were further confirmed by inhibition of dCas9 stably expressed in the 293FT cell line, which was cotransfected with sgRNA, truncated A4b and the EGFP gene for transcriptional activation (Figure S7C). In these confirmational experiments, the proportion of EGFP-expressing cells is a reporter of dCas9 inhibition; it is high when an Acr mutant does not inhibit dCas9, and vice-versa (Figure S7D). In contrast to Δ4, the results showed that the Δ8 mutant efficiently blocked dCas9 activities, similar to FL.A4b (Figure S7E-F). To determine the truncatable residues *in vivo*, we inserted these IDR/IOR-truncated mutants at *attp40*. When they were coexpressed with dCas9_fly, the Δ8 mutants of A4a and A4b were able to rescue the dCas9_fly-mediated phenotypes (Figure 4E and 4F). In addition, all other mutants were as relatively inefficient in inhibiting dCas9 *in vivo*, except Δ4 (Figure 4F). These results demonstrate that certain residues in IDR2 and IOR can be truncated without disrupting A4 function *in vivo*.

Knowing the truncatable residues in IDR2 and IOR of A4 and prototype A4b.LOV_63 photoswitchability in endogenous gene regulation *in vivo* (Figure 3I and 3J), we generated flies expressing “<u>I</u>OR/IDR-<u>T</u>runcated AcrAIIA4-<u>L</u>OV2 Fusion (ITALF)” variants: ITALF1 (∂N64/L86/N87), a partially IDR- and IOR-truncated A4b, and ITALF2 (∂N64-E66/S84-N87), a fully IOR- and partially IDR-truncated A4b (Figure 4G). Alignments of generated alphaFold models of these variants showcase that they are structurally similar to A4b.LOV_63; thus, they occupy the same position in dCas9 as the later (Figure 4H). Comparison of phenotypes showed that ITALF2 did not stably block dCas9_Fly in dark *in vivo* (Figure 4I). In term of photoswitchability, although the two ITALFs were more light-responsive than A4b.LOV_63, ITALF2 was significantly more sensitive to blue light than that of ITALF1 (Figure 4J).

Further, we generated additional A4b-LOV2 variants: ITALF3 with truncations of two IDR2 (∂N64/Q65) and three IOR residues (∂E85-N87), ITALF4 with an entirely truncated IOR, and ITALF5 with a truncation IOR and beyond (∂L82-N87) (Figure 4K). Notably, strong light-response was yielded with ITALF3; truncation of the entire IOR (in ITALF4) led to efficient dCas9_fly inhibition in dark and fair, but not optimal, photoswitchability; truncation beyond the computed IOR (in ITALF5) abrogated the inhibitory activity of dCas9_fly in dark (Figure 4L, Figure S8A). Similarly, IOR truncation of A4a.LOV_63 (ITALF6 and ITALF7) also effectuated reliable dCas9 inhibition in dark and moderate photoactivation under blue light (Figure S8B-D). These findings demonstrate that fine-tuning IOR and IDR length significantly enhances the photoswitchability of the A4b.LOV_63 prototype *in vivo*.

### 5. Optimization of LOV2 and light properties enables rapid photoactivation of ITALF

To further improve *in vivo* functionality, we explored truncation of LOV2 termini, a modification known to enhance photoswitchability (39, 40). Established on that, we designed ITALF3.1 and ITALF3.2 upgraded versions with termini-truncated LOV2 (∂A2/E142) and the reported short-LOV2 (∂L1-T4/K141-L143) (39, 40), respectively (Figure 5A). When coexpressed with dCas9_fly, ITALF3.1 exhibited a marginally high photoswitchability than ITALF3; ITALF3.2 hardly inhibited dCas9 in dark (Figure 5B). Although the short-LOV2 (ITALF3.2) was used to control nanobodies and transcription factors (39, 40), it seems not suitable for AcrIIA4 (Figure 5B). Our data suggest that the hydrophobic leucine residues at the LOV2 termini are critical for maintaining a fully closed LOV2 dark state.

**Figure 5:**
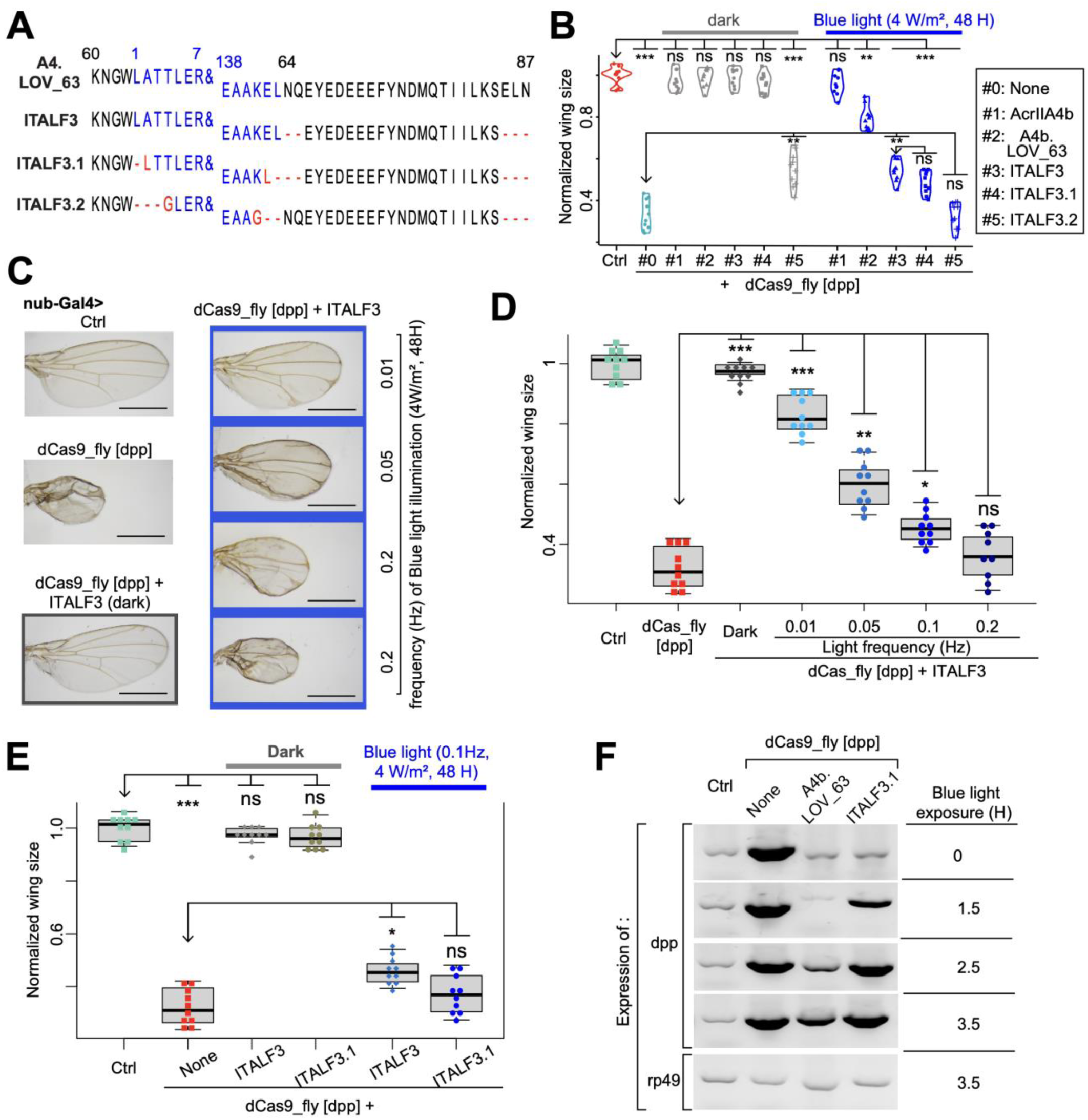
Mutation of LOV2 and adjustment of light properties of ITALF3. (**A**) Peptide sequences of ITALF3, ITALF3.1 and ITALF3.2, whose LOV2 (in blue) of which are truncated, and aligned with A4b.LOV_63 and ITALF3. The ampersand signs (&) are 8 – 137 residues of LOV2. The dash and letter in red represent truncated and mutated residues. (**B**) Plot of normalized wing sizes from dCas9_fly drosophila expressing Acr-LOV2 complexes in **A**. It shows that ITALF3.1 is marginally light-sensitive than ITALF3, and ITALF3.2 is dysfunctional. The larvae of these flies were kept in dark or irradiated with blue light (∼0.08Hz, 4mW/m^2^ for 48H). (**C-D**) Representative images (**C**) and statistical data (**D**) of comparison of wing phenotypes from ITALF3 exposed at different light frequency at a larval stage. The wings were from fly adults, the larvae of which were kept in dark or irradiated with blue light (4mW/m^2^ for 48H). (**E**) Comparison of ITALF3 and ITALF3.1: statistics of wing phenotypes from dCas9_fly drosophila, expressing ITALF3 or ITALF3.1, and the larvae of which were exposed to blue light (0.1Hz, 4W/m^2^, 48H). (**F**). In **B**, **D** and **E**, the labels *ns*, *, **, *** are *P* values not significant, <0.01, <1e-3 and <1e-5, respectively, that were calculated by ANOVA-test of the difference between the normalized wing sizes or eye surface area of control flies and every group.

To optimize the photoswitchability of this system *in vivo*, we also refined light exposure conditions. Compared to that of 24 and 72 hours, 48 hours of light exposure induced reasonable phenotype and mitigated phototoxicity (Figure S9A and S9B). Noting that the low irradiances caused no significant effect to the wings while the high one (10 W/m^2^) was toxic, 4 W/m^2^ irradiance, which is almost bright as for *in vitro* experiments (11, 12), showed reasonable photo-toxicity and light-response in the context of ITALF3 (Figure S9C and S9D). Next, by calibrating the light pulse rate, we remarked a limited degree of phenotypic changes for the low frequency (∼0.01Hz), which is sufficient to maintain adduct state of *Avena sativa* LOV2 that we used (41) (Figure 5C and 5D), unlike other LOV variants (42, 43). Besides, we observed phenotypic changes of dCas9_fly-ITALF3 and of dCas9_fly-ITALF3.1 comparable to those of dCas9_fly alone for the pulse of 0.2Hz and 0.1Hz (4 W/m^2^ and 48 hours), respectively (Figure 5D, 5E and S9E). These indicated that an optimized condition has been achieved for photoswitchability of ITALF3 and ITALC3.1, the slightly better one, in gene regulation *in vivo*. A closer examination at mRNA level show that the maximal upregulation can be reach after 2.5 hours of blue light exposure for ITALF3.1, while over 3.5 hours for A4b.LOV_63 (Figure 5F). In that sense, ITALF3.1, which responds to light significantly faster than A4b.LOV_63, is most probably more light sensitive than the recently developed optogenetic tool, which can reach its maximal upregulation of transgenic genes after 3 hours of light exposure (44). Together, ITALC3.1 achieves an optimally light-sensitive and rapid EnGup *in vivo*.

### 6. ITALF enables investigation of biological functions genes in different tissues *in vivo*

We next sought to demonstrate the application of our light-switchable gene regulation module (ITALF3.1) in optogenetically controlling gene expression of other tissues *in vivo*. We started with investing whether, in adipose tissues, ITALF3.1 can control EnGup such as of *Wnt4* gene, which has been speculated to be involved in autophagy (45). For this investigation of ITALF3.1 in adipose tissues together with the involvement of *Wnt4* in autophagy, we coexpressed ITALF3.1, *Wnt4* EnGup and mCherry-marked autophagy-related gene 8a (Atg8a) using Cg-Gal4 driver. Then, before mounting their adipose tissues on slides, larvae were subjected to different experimental paradigms: (#1) fed normally; (#2) starved for 6 hours in the dark; (#3) starved for 6 hours, during the first half (3 H) of which the larvae were exposed to blue light; and (#4) starved for 6 hours, during the last half of which the larvae were exposed to blue light (Figure 6A). Atg8a is expressed in nucleus under normal condition, and forms puncta in cytoplasm when larvae are under stress or starved (Figure 6B), as previously reported (46). But surprisingly, the *Wnt4* EnGup suppresses the formation of these Atg8a puncta; meaning that our results provide insights of the involvement of *Wnt4* in autophagy that previously speculated (45). More intriguingly, the coexpression of *Wnt4* EnGup and ITALF3.1 under food stress allowed Atg8a-puncta formation in the experimental paradigms #2 (in the dark) and #4, and suppression of this formation in #3 (Figure 6B and 6C). Thus, our system enables to optogenetically control biological process in other tissue as such the *Wnt4* EnGup-mediated autophagy suppression in adipose tissue.

**Figure 6:**
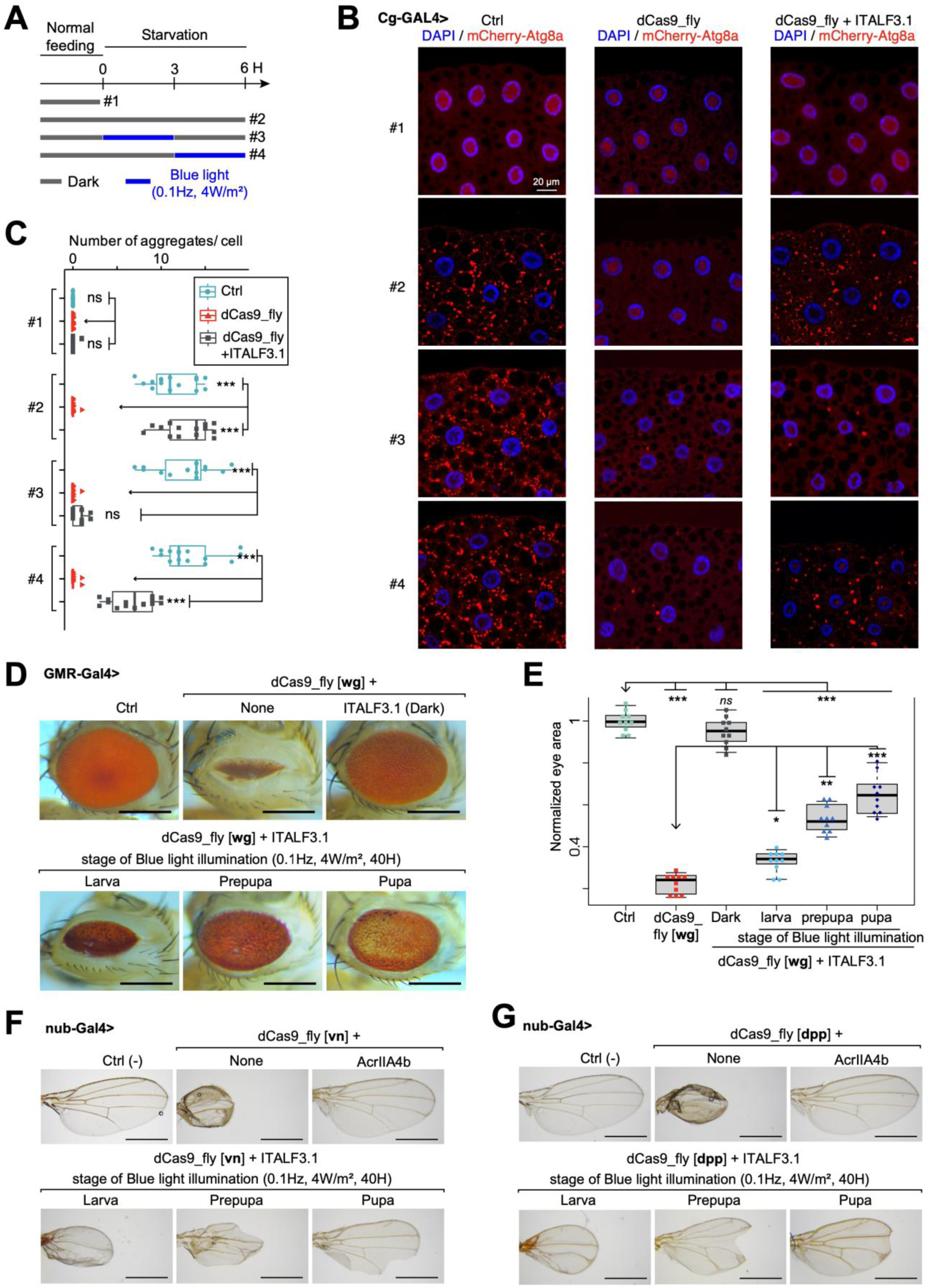
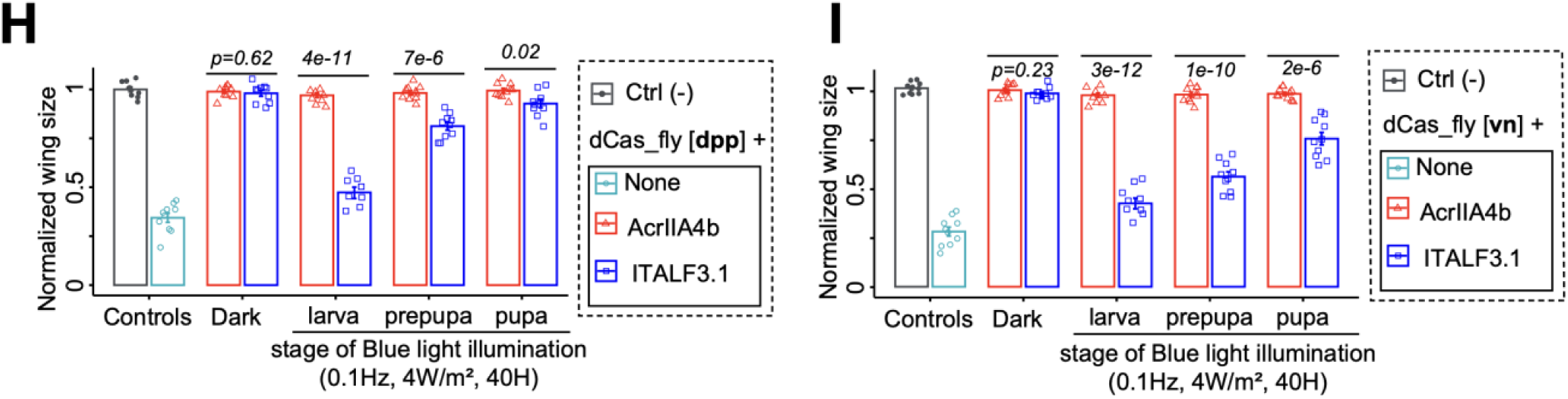
Application of ITALF on targeting different tissues and genes *in vivo*. (**A**) Experimental paradigm for investigating ITALF3.1-mediated *Wnt4* EnGup through expression of mCherry-Atg8a (in red). Larvae were normally fed or starved either in the dark (dark gray lines) or under a blue light (blue lines), before mounting their adipose tissues. (**B**) Confocal images of L3 fat body adipocytes showing localization and expression of mCherry-Atg8a in red (driven by Cg-GAL4). Nuclei stained with DAPI (blue). (**C**) Plot quantifying mCherry-Atg8a puncta per cell from data in **B**. (**D**) Image samples illustrate the eye phenotypes of dCas9_fly drosophila adults expressing ITALF3.1. These flies, whose larvae, prepupae or pupae were illuminated under a blue light pulse (0.1Hz, 4mW/m^2^ for 40H), transcriptionally activated *wg* gene. The scale bars are 250 μm. (**E**) Plot summarizes the data in **D**. (**F-G**) Representative images of wing phenotypes from dCas9_fly flies, expressing ITALF3.1 or AcrIIA4b and dCas9_fly targeting *vn* gene in **F** or *dpp* in **G**. The larvae, prepupa or pupae of these flies were irradiated with blue light (0.1Hz, 0.4mW/cm^2^) for 40H. The scale bars are 500 μm. (**H-I**) statistics from **F** in (**H**) and **G** in (**I**). *P* values calculated by ANOVA-test are the mean of normalized wing sizes from dCas9_fly flies expressing ITALF3.1 compared to those expressing AcrIIA4b. In **C** and **E**, the labels *ns*, *, **, *** are *P* values not significant, <0.01, <1e-3 and <1e-5, respectively, that were calculated by ANOVA-test of the difference between the normalized wing sizes or eye surface area of control flies and every group.

To confirm this ability to target any tissue, we generated a fly line for wingless (*wg*) EnGup, leading to robustly smaller eyes – compared with *dpp* and *vn* EnGups – when driven by GMR-Gal4 and also smaller wings when driven by nub-Gal4 (Figure S10A-S10D). Capitalizing on these results, we temporarily upregulated *wg* expression in eyes by coexpression of dCas9_fly and ITALF3.1 at three different developmental stages. Light-irradiated larvae gave rise to adults with narrower eye area, whereas the pupae to flies with facetless eye area (Figure 5D). The dark-light difference of eye phenotypes was significant at every stage, although that at the larval stage was most significant (Figure 6E). Our results using our light-switchable gene regulation module show the ability of the *wg* gene to disrupt photoreceptor formation when expressed during *Drosophila* early developmental stages (Figure S10E), similar to what was previously shown by exogenous gene overexpression (47, 48). These, therefore, validate that our light-switchable module can be used for any tissue and at any developmental stage.

To generalize this approach, we also used our light-switchable system to regulate *vn*, *dpp* and *wg* expression in wings. When *vn* upregulation was activated by light at larval and prepupal or pupal stages of development, we observed, respectively, proximally distorted and distally deformed wings in fly adults (Figure 6F). Light-activation of *dpp* at larval and prepupal or pupal stages caused, respectively, proximally distorted and distally notched wings (Figure 6G). Light-activation of *wg*, finally, marginally reduced wing size when expression was turned on at larval and prepupal but not pupal stage (Figure S10F and S10G). The upregulation of *vn* and *dpp* both significantly reduced adult wing size (Figure 6H and 6I), in agreement with previous studies using exogenous gene overexpression (49–52). In summary, our results prove the efficiency of our system to optogenetically regulate any endogenous gene expression.

## DISCUSSION

This study represents the first report of AcrIIA4 and AcrIIA2 application in *Drosophila* and, more broadly, in animal models. Until now, the use of these Acr families – well-known CRISPR antagonists – *in vivo* has been restricted to yeast (53). In *Drosophila*, an inhibitor or antagonist is needed to defend against toxicity of continuous CRISPR-Cas activity, which is lethal when expressed in vital tissues (7). Besides preventing CRISPR from off-target effects (54) and cellular toxicity (55, 56), Acr proteins have been demonstrated to maintain on-target editing while reducing off-target editing in human cells (32), to generate “write-protected” cells that prevent future gene editing as well as to control CRISPR-based gene regulation circuits (57). Here, we demonstrate for the first time that wild-type AcrIIA2, when sufficiently titrated *in vivo* at room temperature, efficiently inhibits Cas9 (Figure 3H-I, Figure S6), expanding the functional repertoire of Acr proteins.

Likewise, we introduce a CRISPR-based optogenetic system to modulate gene expression in *Drosophila*. Here, we demonstrated the application of ITALF3.1 to phenotypic modelling and autophagy investigation, but users can combine this system with Cas9, CRISPRi (8) and dCas9_fly (7) – through already available stocks – to mediate light-dependent editing and expression of any endogenous gene. This capability allows for *in vivo* cellular reprogramming and disease modeling. Moreover, the versatility of ITALF in targeting genome-wide genes complements the presently available optogenetic approaches designed for transgenes (4–6). Conceptually, despite the fact that LOV2-inserted Acr proteins have been used in cell cultures (25, 26) and in *E.coli* (27), the current work is the first to develop such system in an animal. Notably, the success of prototype A4b.LOV_63 was not optimally predicatble from previous findings, carried out *in vitro* cell cultures, which emphasizes that the insertion of LOV2 around the IDR E66 or Y67 is preferable for AcrIIA4 (26). This cell-culture-and-in-vivo difference could be due to: i) the residue mismatches of AcrIIA4 we used; ii) or most plausibly, the strong binding specificity of IDRs of AcrIIA4 in inhibiting Cas9 *in vivo*. Such binding specificity of IDRs, which instigates more in-depth investigation of Acr-Cas9 interactions, has been pointed out to determine the functionality of transcriptional factors in yeast and mouse (10–15).

We optimized our prototype by refining the length of IDR and IOR of AcrIIA4 (Figure 4, Figure S9). While IDR is a widely accepted concept, its prediction in proteins remains an ongoing challenge, with computational methods continuously evolving (22, 58–61). The generality of IDRs in functional interactions is still being explored (58), and their presence in small AFDB proteins has not been previously reported. Our simple approach provides another angle for understanding IDRs of small proteins. Regardless of the size of Acr proteins, our data support the ability of IDRs to regulate the stability and function of proteins as reported in other studies in *Drosophila* (62–64). On the other hand, IOR is a newly proposed concept (21). The IOR of AcrIIA4a is more acidic and marginally longer IOR than that of AcrIIA4b, a reason why A4b.LOV_63 is more light-responsive compared to the other prototypes (Figure 3I-J). Likewise, IOR-truncated ITALF4 and ITALF6 along with ITALF7 are more light-sensitive than the A4b.LOV_63 and A4a.LOV_63, respectively. These imply that residues of IOR in AcrIIA4, and probably in small-sized proteins, can be mutable and truncatable for a particular purpose. Structurally, IOR of AcrIIA4 does not more strongly interface Cas9 than IDRs; considering this, we speculate that IOR mechanistically contributes to Acr-CRISPR interaction as a kinetic stabilizer of Acr functionality, which may have important implications for protein engineering.

In summary, this study pioneers the use of Acr proteins *in vivo* and their engineering for light-regulated gene expression in *Drosophila*. These first prototypical optogenetic tools were optimized by truncating residues of IDR and IOR, which are structures herein characterized for the first time *in vivo* for small sized Acr proteins. The broader applicability of IDR and IOR concepts in other small proteins remains an open question for future research.

## MATERIAL AND METHODS

### Plasmid construction and cloning

All constructs were generated with NEBuilder HiFi DNA Assembly Cloning Kit (NEB, #E5520S) or 2× Seamless Cloning Mix (Beyotime, #D7010M). Briefly, DNA sequences synthesized by Xiang Hong (xianghongbio.com) or available plasmids from Addgene were amplified according to 2 × Taq Plus Master Mix II protocol (Vazyme, #P213), and used as templates; pValium-ROE10 (65), pScript-Renilla (66), and lentiCRISPR v2 (Addgene, #52961) were linearized, with KOD One^™^ PCR (Toyobo, #KMM-101), and used as vector backbones for plasmids to be injected in *Drosophila*, expressed transiently and expressed stably in cells, respectively. The fragments and vectors were purified by V-ELITE Gel Mini Purification kit (ZOMANBIO, #ZPV202) and assembled by 2× NEBuilder Hifi Master Mix (NEB, #E2621s) or 2× Seamless Cloning Mix Kit (Beyotime, #D7020S).

### sgRNA design

Single guide RNAs (sgRNAs) for upregulation of *wg* gene (ID: FBgn0284084) were designed using the *Drosophila* Resource Screening Center Find CRISPRs v2 online tool (www.flyrnai.org/crispr/). Three pairs of sgRNAs, which were additionally screened for potential off-targets with cas-OFFinder (http://www.rgenome.net/cas-offinder/) and targeted 300 to 800 bp upstream of *wg* gene, were selected and synthesized. Each pair of oligo DNA was cloned in flySAM construct according to the protocol (67).

### *Drosophila* stocks and fly husbandry

All *Drosophila* stocks were maintained on standard cornmeal/sugar/agar medium (7). Experiments were conducted at 25°C. *Drosophila melanogaster* stocks used in this study are listed in Table S2. Transgenic flies were generated by the injection of plasmids with UAS-driven Acr or Acr-LOV2 fusion proteins and dCas9_fly for *wg* EnGup into *y sc v nanos-integrase; attP40* or *y sc v nanos-integrase;; attP2*.

### Cell culture and stable cell line generation

Human embryonic kidney 293T (HEK293T) cells were cultured in DMEM (Gibco, #C11995500BT) supplemented with 10% (vol/vol) fetal bovine serum (FBS; Gibco, #16000-044) and 1% (vol/vol) penicillin solution (Beyotime, #ST488-1) at 5% CO2 and 37 °C in a humidified incubator. The cells were PCR tested for mycoplasma contamination, and passaged every 2–4 days.

For stable cell lines, Cas9 and dCas9-3(VP64) cloned in lentiCRISPR v2 vector plasmids were injected into HEK293T cells according to the established protocol (68). Briefly, mixture of 5 µg of each of these lentiviral plasmids, 2.5 µg of pMD2.G (addgenes, #12259), 3.75 µg of psPAX2 (addgenes, #12260), and 12.5 µL of Lipofectamine 3000 (Invitrogen, #L3000015) in OptiMEM (Life Technologies) was delivered into HEK293T cells in a 60-mm dish (Nest, #706001) at 80-90% confluency. After 6 hours, the culture medium was changed to DMEM with 10 % FBS, then added with 4µg/mL of blasticidin for selection 24 hours later. Passaging and selection were carried out twice more, before storing the stable cell lines at -80°C in M5 Hiper medium (Mei5Bio, #CS699-01).

### Plasmid cotransfection

For transient expression, stable cell lines were (co-)transfected employing an optimized polyethyleneimine (PEI)-based protocol. Briefly, 5 × 10^4^ cells cells were seeded into a 12-well plate, cultivated overnight to reach 60-70% confluency, and cotransfected with 2 μg/well of a plasmid mixture (1:1 sgRNA&Acr/reporter plasmid ratio). The plasmid mixture was diluted in 50 μl OptiMEM, subsequently mixed with 50 μl PEI solution containing 3.2 μl of PEI (1 mg/ml in ddH_2_O; Polysciences, #24765-2) and 46.8 μl OptiMEM, incubated for 15 min at 25°C, and dropwisely added to the cell culture. The culture medium was renewed 6 hours after transfection.

### Computational modeling

Although the Acr proteins we used were available from alphaFold database (36), we performed the modeling of Acr-LOV2 fusion proteins and Acr-dCas complexes using locally-installed version of the AlphaFold_2.0_multimer.v3 and more recent AlphaFold3 (https://github.com/google-deepmind/alphafold3), along with ColabFold from Github (https://github.com/YoshitakaMo/localcolabfold) and the environmental databases (https://colabfold.mmseqs.com). We compared the top five ranked outputs and selected Rank 1 to prepare the figures, which were rendered, aligned with SpyCas9–sgRNA–AcrIIA4 (PDB 5VW1) as reference, with the UCSF ChimeraX-1.7.1 software (https://www.cgl.ucsf.edu/chimerax/).

### Structural analysis

The per-residue local distance difference test (pLDDT) confidence scores of Acr proteins downloaded from the AlphaFold (36) were retrieved from the atomic coordinates of a protein residue in a pdb file. These coordinates were mapped in 2D using bio3D library in RStudio.

Electrostatic potential charges and hydrophobicity of surfaces of AcrIIA4 pdb files were rendered by ChimeraX. However, their exact value from each residue was extracted using surface_analysis library (https://github.com/liedllab/surface_analyses). As for the local charge distribution of proteins, unlike the other structural analyses of 3D structured protein, it was extracted from a primary structure. The average charge of each residue was computed with seqinr and bio3d libraries on RStudio.

### Contact map analysis

Acr-dCas9 complex pdb file was further processed with Jupyter Notebook (version 6.5.4) with nglview library. The distance matrices between Acr and dCas9 chains and between two residues were calculated using biopython and numpy (version==1.23.4) packages, and those with distances below the 8Å threshold were defined as in contacts. Then, rpy2 and tzlocal packages were employed to 2D-map these distances. Then, the contact map was refined by removing the residues of dCas9 with distances above the 10Å.

### Luciferase assay

The luciferase assay was carried out 2–3 days post-transfection. Cells cultivated in 12 well plates, the cultured media of which were aspirated, were washed with 1xDPBS (Corning, #21-031-CV) lysed with 100µL/well of cell lysis buffer (Beyotime, #RG027-1) and incubated on ice for 10 min. Then, Firefly Luciferase activities were measured according to the previously adapted protocol (69). Briefly, 50 µL of cell lysates were transferred to one well of 96-well white opti-plate (PerkinElmer #6005290), then mixed with 100µL Bio-Lite Luciferase Assay Substrate (Vazyme, #DD1202) that had been dissolved with its buffer. The opti-plate, after incubation at room temperature for 10 minutes, was loaded into GloMax-Multi Microplate Reader (Promega), which was set for steady-Glo to measure the luminescence for 2 sec. All results were normalized with a positive control that was loaded in the first column of the opti-plate.

### FACS analysis

Forty-eight hours after transfection, the medium was removed, cells were detached from the each well using trypsin EDTA, transferred into 1.5 mL and centrifuged at 300g for 7 min at 4 °C. The supernatant was discarded, and the cells were resuspended in 1xPBS. The fluorescence-activated cell sorting analyses were carried out using FongCyte Series 3-Laser 14-Color (Chellenbio, Beijing Challen Biotechnology Co., Ltd) with the settings: 450 nm laser and a 500/50 filter with a photomultiplier tube (PMT) set at 275 V. For each sample, 10^5^ cell events were collected.

### Blue light assay

Adult flies were kept in rearing conditions in the dark: in a box placed in an incubator at 25 °C with humidity of 60–65%. Every parental population was raised in the same conditions, and transferred within 24 hours into a fresh vial filled with standard fly food (7). Offspring, 3–4^th^ instar larvae or pupae, from these crosses were shortly screened under a microscope, and up to 35 of the correct genotypes were transferred into a 6 cm dish (NEST, #706001), containing 1–2mm layer of semi-transparent fly food (50% standard fly food media (7), 40% apple juice, 1% agar, 9% water; boiled and cooled down to 70°C before use). The lid-covered and sellotape-sealed dishes were placed either in dark or in a handmade blue light box – a box painstakingly covered with aluminum foil from inside and illuminated with a custom-made 450–460 nm blue light (Xuzhou Ai Jia electronic technology Co.LTD), which was powered by JUNCTEK JDS8080 80MHz Signal Generator (https://www.junteks.com/). After treatment (for 30 to 48 hours), the offspring were transferred into a fresh vial and put back into the dark conditions.

### Phenotypic analysis

Female adult wings (12–15 per genotype) were dissected, rinsed with ethanol, mounted in a solution of 1:1 acetone to Permount (Thermo Fisher Scientific) on a glass slide, and imaged on Zeiss Axioskop 2 microscope. Representative images were shown for each genotype, but the characteristics (size, length, width, …) of 10 wings were extracted via python scripts on Spider environment (version 5.4.3) using OpenCV, Numpy and Skimage libraries. As for eye phenotypes, CO2-anaesthesized 20 female adults were plated on a ∼3 mm layer of 0.4% agar (boiled and cooled down to 50°C before use); eye images were taken with a Zeiss Axioskop 2 microscope, and processed with ImageJ (version 2.14.0).

### Confocal imaging

The adipose tissues were dissected from 3^rd^ instar larvae with fine-tip forceps, washed in PBS for 2 × 15 min, and mounted on a glass slide with a drop of DAPIVectashield (Vector Laboratories, #H-1200). The mounted tissues were observed under a Nikon AXR NSPARC using the 63×/1.42 NA oil objective or ZEISS LSM780 confocal microscope equipped with an objective 63× oil Plan-Apochromat (NA 1.4). The images were acquired with laser lines at 488 and 640 nm, and processed with NIS-Elements software (Nikon) and python scripts.

### Viability analysis

Newly hatched adults from each fly line were cultured in an incubator at 25 °C with humidity of 60– 65% and a 12-h dark/light cycle. They were transferred every day into new vials with fresh medium to prevent non-age-associated mortality like accidental trapping on food, and the numbers of dead flies were recorded. These procedures were continued for 75 days or until all flies were dead. The Kaplan– Meier method was used to present survival data and the log-rank test was used to compare survival distributions.

### Quantitative RT-PCR

Total RNA was isolated from early hatched adults, 50 wings per genotype were dissected in Schneider’s medium (Sigma Aldrich). The Quick-RNA Microprep kit (Zymo Research, #R1050) was used following the manufacturer’s instructions to isolate RNA, then it was incubated with DNase I (Promega, #Z358A) at 37°C for 30 minutes and treated with the RNA Clean and Concentrator-5 kit (Zymo Research, #R1015). A total of 1 μg of RNA was used as a template for cDNA synthesis using Moloney Murine Leukemia Virus reverse transcriptase (M-MLV RT) (Invitrogen, #28025013). Results were normalized against rp49 expression.

### Statistical analysis

For all experiments, bars indicate means, individual data points correspond to individual biological replicates. For the *in vitro* experiments, all data represent the mean ± s.d. of three independent experiments (*n* = 3), with three replicated for the luciferase experiments. For animal experiments, each genotype or treatment consisted of ten female *Drosophila* (*n* = 10) that displayed representative phenotypes. All group comparisons were analysed by one way ANOVA-test (cut-off *P* < 0.01) using R script on Rstudio. No blinding was performed in all experiments.

## Data availability

All data supporting the findings of this study are available within the paper and the supplementary Information. Plasmids and relevant fly lines are also available, and will be stored in public data base.

## ACKNOWLEDGMENTS

We very much appreciate the help of Jian-Quan Nís lab (School of Medicine, Tsinghua University) and the Tsinghua Fly Center (THFC) for the generation of all transgenic flies. We acknowledge the contribution of Ke Yang and Yuwei Huang in José Pastor-Parejás lab for technical assistance and replication of key experiments. We are very grateful for the internal reviewers of Tsinghua University specially Prof. Jun-Jie Liu, Prof. Wei Zhang and Prof. Xin Liang. We also thank Wei Zhanǵs lab for material and technical support. We express our gratitude to all members of Yichang Jiás group, specially to Xichen Zhou, Dr. Xue Zhang, Dr. Liang Guo for reproducing experimental key findings, providing plasmids and for their support and critical feedback on the manuscript. The help from Angella Zhao (Neurosciences, Rice University, USA), Cui Wenjia (Peking University), and Jessica Ye and Peggy Chan (Trainees in Yichang Jia’s lab) for fly experiments and plasmid construction are also appreciated. For the imaging and flow cytometry, we sincerely thank Dr. Xiaohe Li (Anming Menǵs lab, Life Sciences, Tsinghua University) and Rong Wu (Life Sciences, Tsinghua University), respectively.

