## Supplementary Figures for "Optimized optogenetic anti-CRISPR for endogenous gene regulation in *Drosophila*"

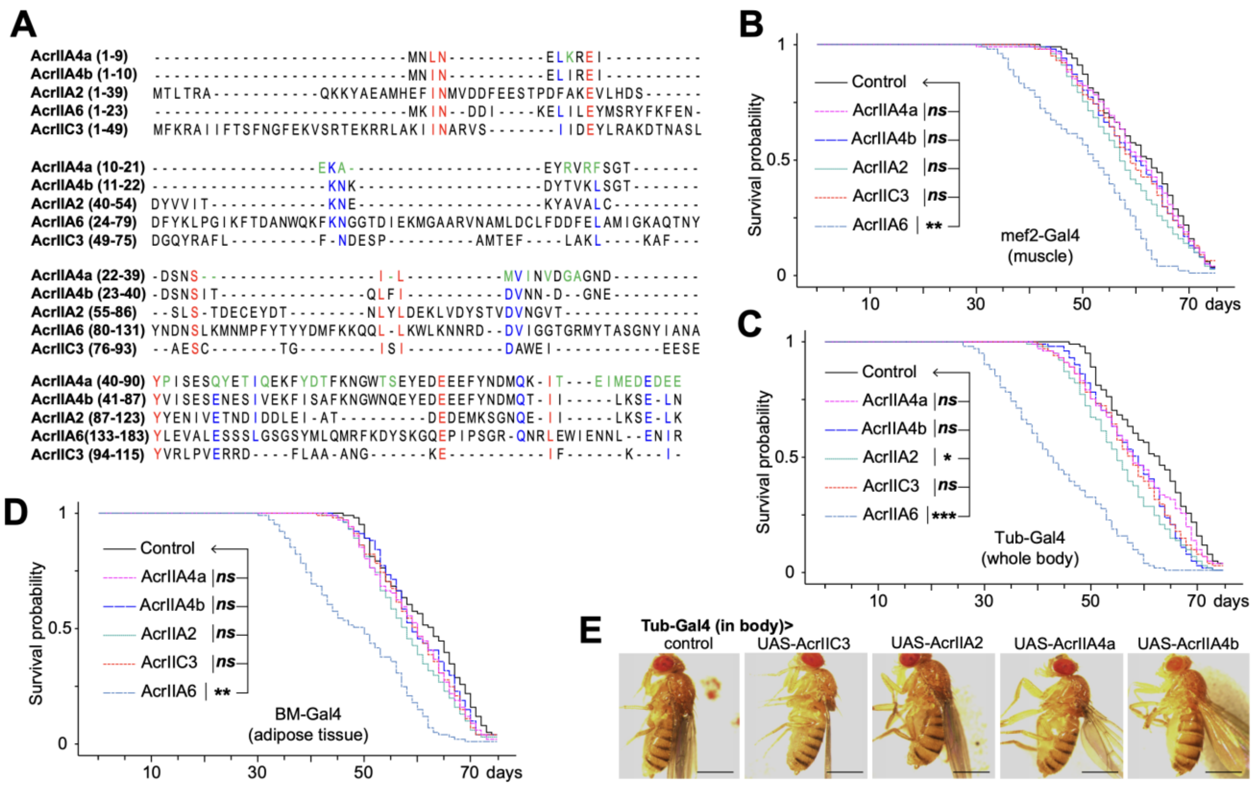


**Figure S1. Acr proteins and their toxicity in *Drosophila*.** (**A**) Multiple sequence alignment of five Acr proteins. The red characters are the conserved residues in all sequences, the blue in four sequences, and the green are the residues differentiate AcrIIA4a from AcrIIA4b. (**B-D**) Survival curves of flies expressing each Acr at different tissues: muscle (**B**), whole body expressing tubulin (**C**), and fat body (**D**). P values *ns*, *, ** and *** are respectively not significant, <0.01, <1e-3 and <1e-5 compared to the normalized control phenotypes by Log-rank test with n = 4 x 25 individual flies. (**E**) Physiology of *Drosophila* generally expressing each potentially non-toxic Acr protein. The scale bars are 500 µm.

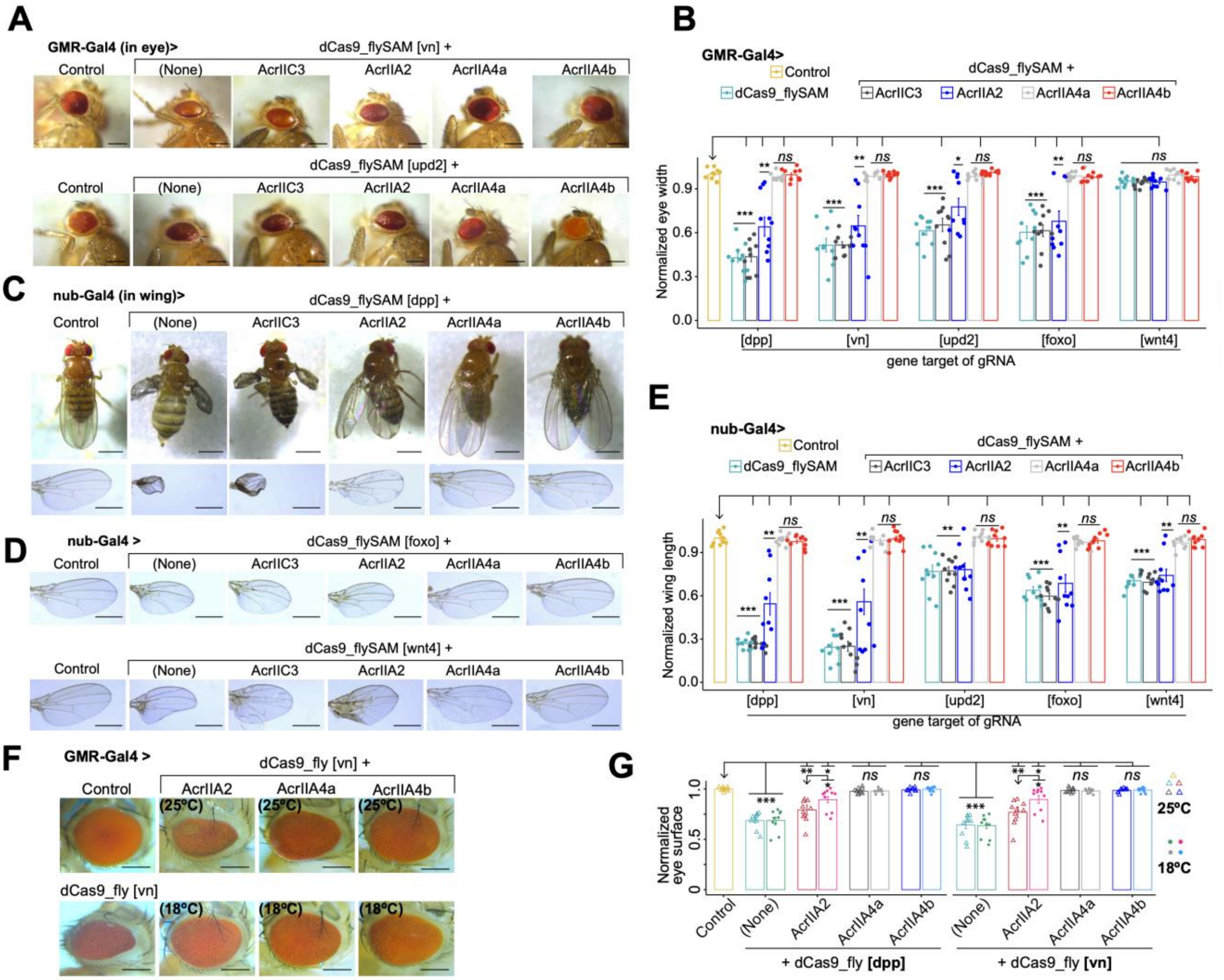

**Figure S2. AcrIIA4a&b effectively inhibit dCas9_fly.** (**A**) Phenotypical eyes of dCas9_fly-targeted *vn* and *udp2* genes, coexpressed with each Acr protein. (**B**) Histogram summarizes the eye width patterns of extended data in **A**. (**C, D**) Nub-Gal4 driven wing phenotypes of Acr-coexpressed dCas9_fly for *dpp*, *upd2* and *wnt4* genes. (**E**) Graph of wing length recapitulates **C** and **D**. (**F-G**) Eye surface representatives (H) and summarized data (I) from flies coexpressing AcrIIA2 or AcrIIA4 and dCas9_flySAM targeting *dpp* and *vn* genes, that were bred in different temperatures. In **B, E** and **G**, every single value is normalized with the mean of normal control. Values are represented as means ± s.d.; n = 10 female flies. *P* values were calculated by ANOVA-test; *ns*, *, **, ***are respectively not significant, <0.01, <1e-3 and <1e-5 compared to that of normal control. The scale bars in **A&F and C&D** are in 250 µm and 500 µm, respectively.

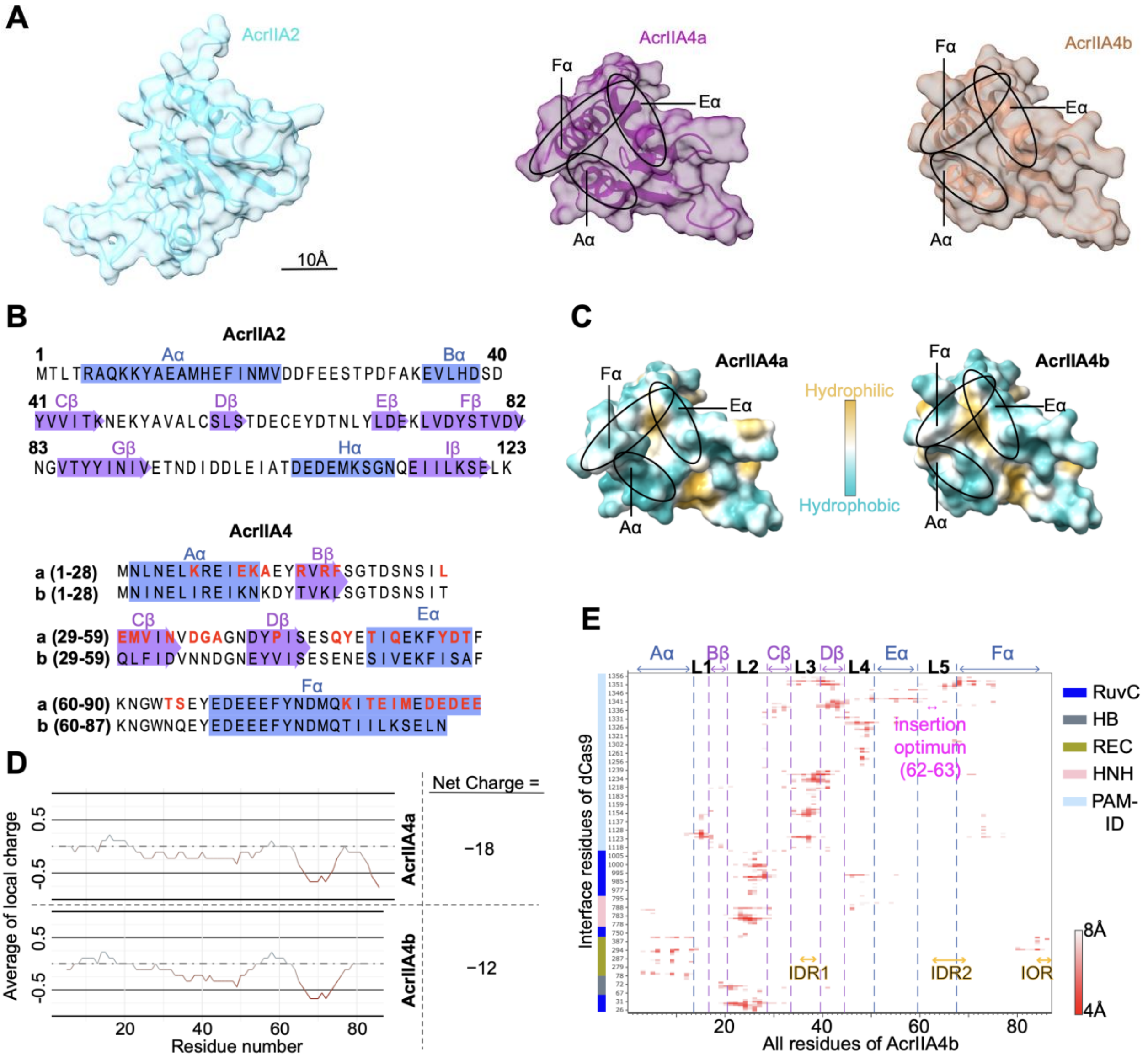


**Figure S3. Further structural analyses and visualization of Acrs.** (**A**) Side views of ribbon representation with semi-transparent surface of AcrIIA2, AcrIIA4a and AcrIIA4b (homology PDB ID 5VW1), colored in cyan, dark pink and sienna, respectively. (**B**) Schematic representations of the secondary structures AcrIIA2 and AcrIIA4a aligned with AcrIIA4b. The amino acid sequences of alpha-chains and beta-sheets are colored in blue and medium orchid, respectively. The red characters are the residues differentiate AcrIIA4a from AcrIIA4b. (**C**) Surface representations with surface hydrophobicity of AcrIIA4a and AcrIIA4b models. (**D**) Local charge distribution of the two AcrIIA4. The charge net of each Acr is indicated. (**F**) Contact map of dCas9-AcrIIA4b complex interfaces within 8Å distance of interactions. Labeled within also are the α-chains, β-sheets and loops (L1-5) of AcrIIA4b. The optimum insertion sites and IDRs along with IOR are spotlighted in magenta and orange, respectively.

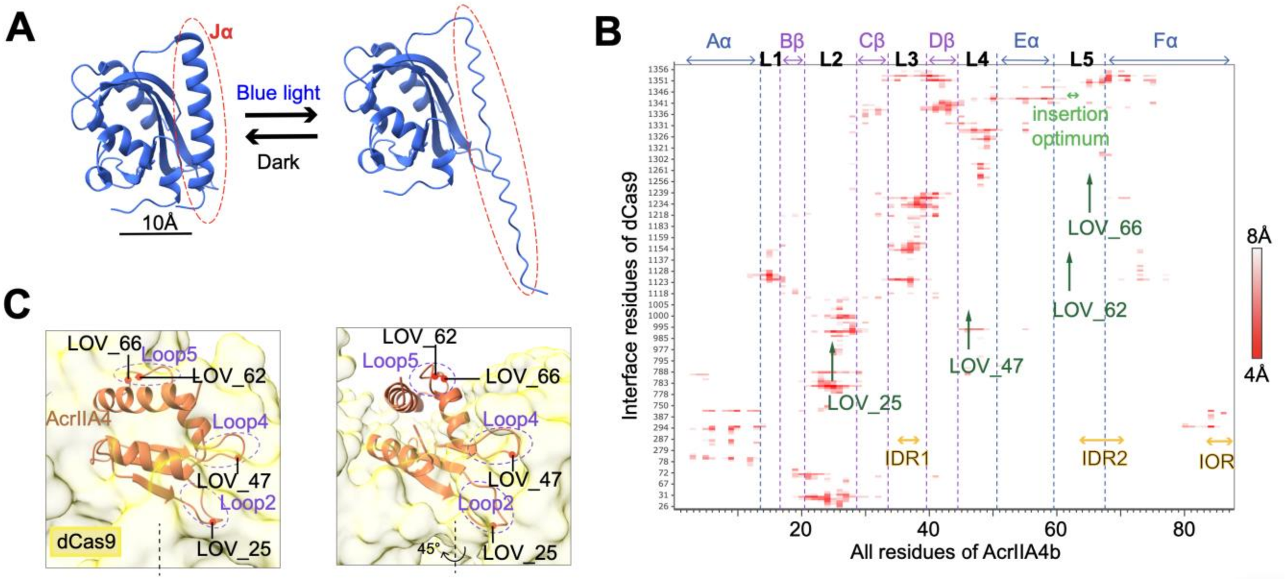


**Figure S4. Selected position for insertion of LOV2.** (**A**) Sides views of LOV2 in dark-state (PDB ID 2V0U) and in light-adapted state. The C- and N-termini are as close as 10Å in the dark, but the Jα helix (dashed red circle) unfolds upon photoactivation. (**B**) Contact maps of predicted the interface residues of dCas9-AcrIIA4b within 8Å distance of interactions and the four selected LOV2-insertion sites (labeled in dark green). The "insertion optimum" label (green) indicate the optimum position for the insertion of LOV2 in AcrIIA4b. The alpha-chains, beta-sheets, loops and IDRs along with IOR are respectively labeled in blue, magenta, black and orange. (**C**) AlphaFold representation of zoomed in views of AcrIIA4b (dark orange) inside of DNA binding pocket of dCas (yellow). It highlights selected insertion sites (25, 47, 62 and 66) of LOV2 in loops (2/4/5; in magenta) of AcrIIA4b. The first view (left) is rotated to 45° angle in the second (right) along the vertical axis (dashed line).

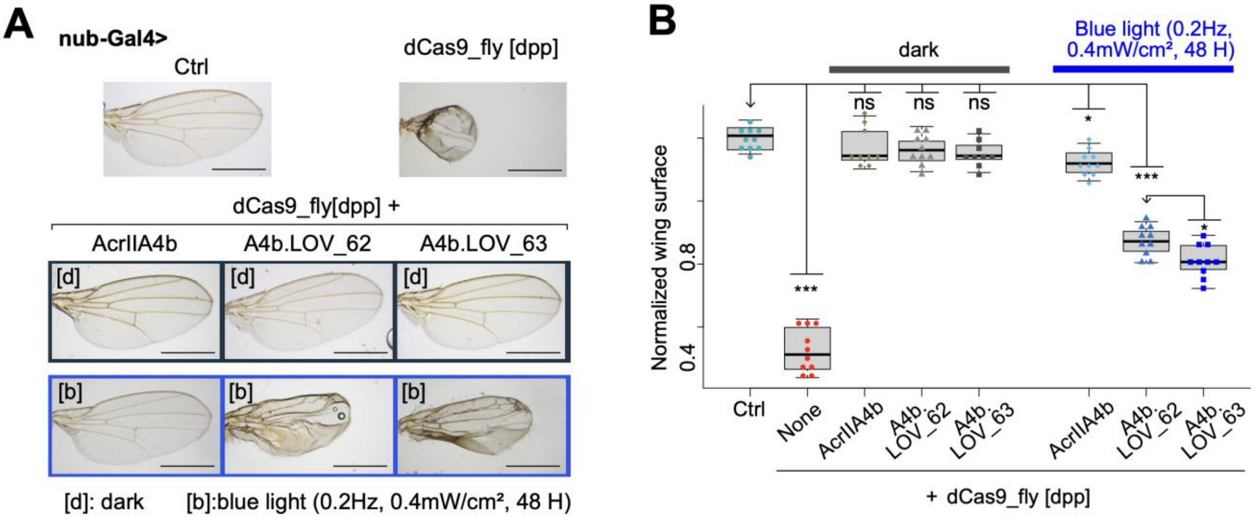


**Figure S5. Comparison of LOV2 insertion in 62 and 63 residues of AcrIIA4 *in vivo*.** (**A**) Images displaying the comparison between the insertion of LOV2 at the positions 62 and 63 of AcrIIA4b. The scale bars are 500 µm. (**B**) Data summary of normalized wing sizes described in **A**. It displays that A4b.LOV_63 is more light-responsive that A4b.LOV_62 for *in vivo* drosophila. The *P* values were calculated by ANOVA-test, with n = 10 female flies; *ns*, *, ** and *** are respectively not significant, <0.01, <1e-3 and <1e-5 between the normalized phenotypes of the arrowhead-pointed group and those of other groups.

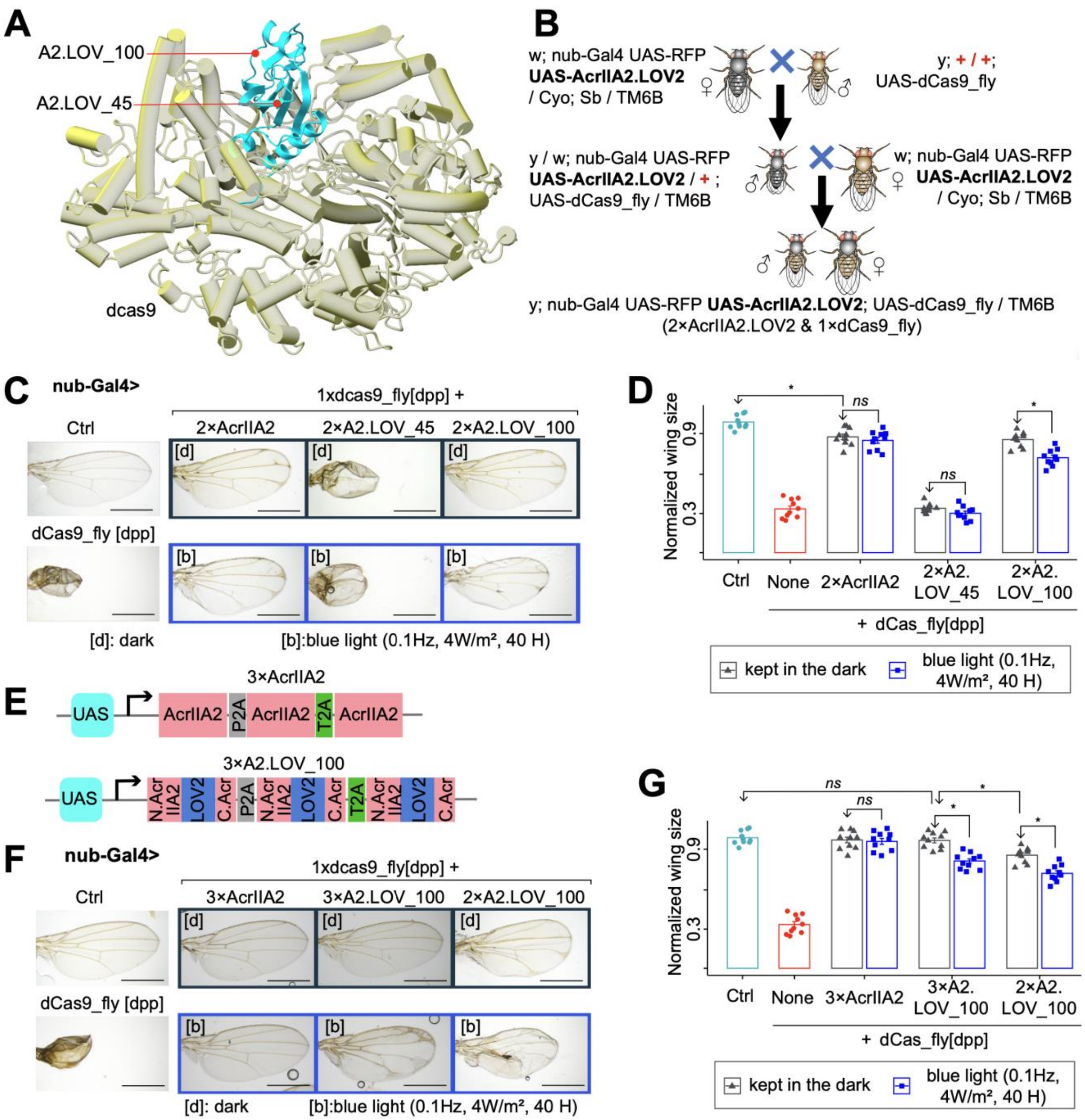


**Figure S6**: **Responses of LOV2-inserted AcrIIA2 to blue light.** (**A**) AlfaFold model dCas9 in complexity with AcrIIA2, highlighting the two selected sites for insertion of LOV2. (**B**) Crossing scheme for generating 2×AcrIIA2.LOV2 coexpressed with 1×dCas9_fly. (**C**) Representative images of wings from adult flies, the larvae of which were lightened under blue light pulse or kept in the dark, expressing dCas9_fly along with variants of AcrIIA2. (**D**) Data summary of normalized wing sizes described in **C**. It emphasizes that 2×AcrIIA2.LOV2_100 is light-responsive. (**E**) Construct for expression of AcrIIA2 and AcrIIA2.LOV2_100 for triple dose expression. (**F**) Illustrative images showing that triple dose of AcrIIA2 can efficiently inhibit dCas9_fly. (**G**) Data summary of normalized wing sizes described in **F**. In **D** and **G,** *P* values were calculated by ANOVA-test with n = 10 female flies; *ns* and * are respectively not significant and <0.01. In **C** and **F** scale bars are 500 μm.

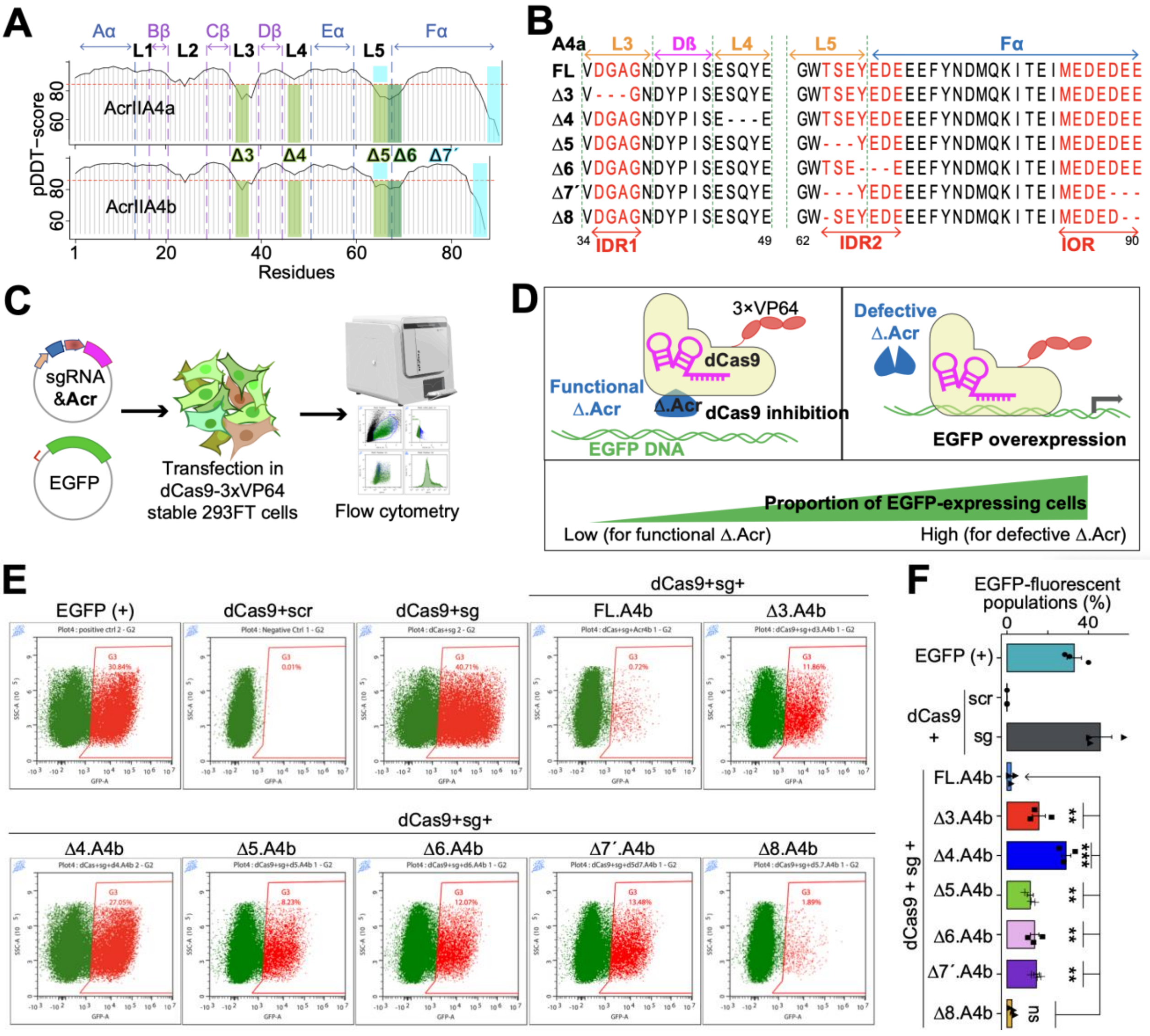


**Figure S7. Identification of truncatable residues in IDRs/IOR of AcrIIA4.** (**A**) pLDDT-scores of AcrIIA4a&b spotlighting their IDRs and IOR for truncation targets colored in yellow green or forest green (Δ3-6) and cyan (Δ7´). (**B**) Partial amino acid sequences of truncated AcrIIA4b variants aligned with the full-length one (FL). Δ7´ indicates a double truncation, and Δ8 is deletion of tree residues in AcrIIA4a (∂64/89/90). The secondary strunctures (alpha-chains, beta-sheets and loops) are outlined. The IRDs and IOR are marked in red characters. The dash (-) represents a single residue deletion. (**C**) Experimental workflow for further investigating the efficiency of truncated Acrs using stable cell line expressing dCas9-3×VP64. Cells stably expressing dCas9-3×VP64 were cotransfected with each truncated Acr, sgRNA targeting 6×crRNA sequence upstream of EGFP, and a reporter EGFP. (**D**) High proportion (in %) of EGFP fluorescence indicates that dCas9 mediates the transcriptional activation of EGFP, while low proportion represents that dCas9 is inhibited by an Acr variant. (**E**) Flow cytometry analysis of stable cell lines transfected with U6-sgRNA, CMV-AcrIIA4b (Full length or truncated) and 6×crRNA-EGFP. The scatter plots are gated on EGFP fluorescence of the negative and positive controls. (**F**) Barplot of statistical data illustrates **E**. Bars represent mean values, error bars the standard deviation and dots individual data points from with n=3 independent experiments. The *p* values * < 0.01, ** <1e-3, *** < 1e-5 were calculated by one-way analysis of variance (ANOVA) between WT and each truncated Acr variant.

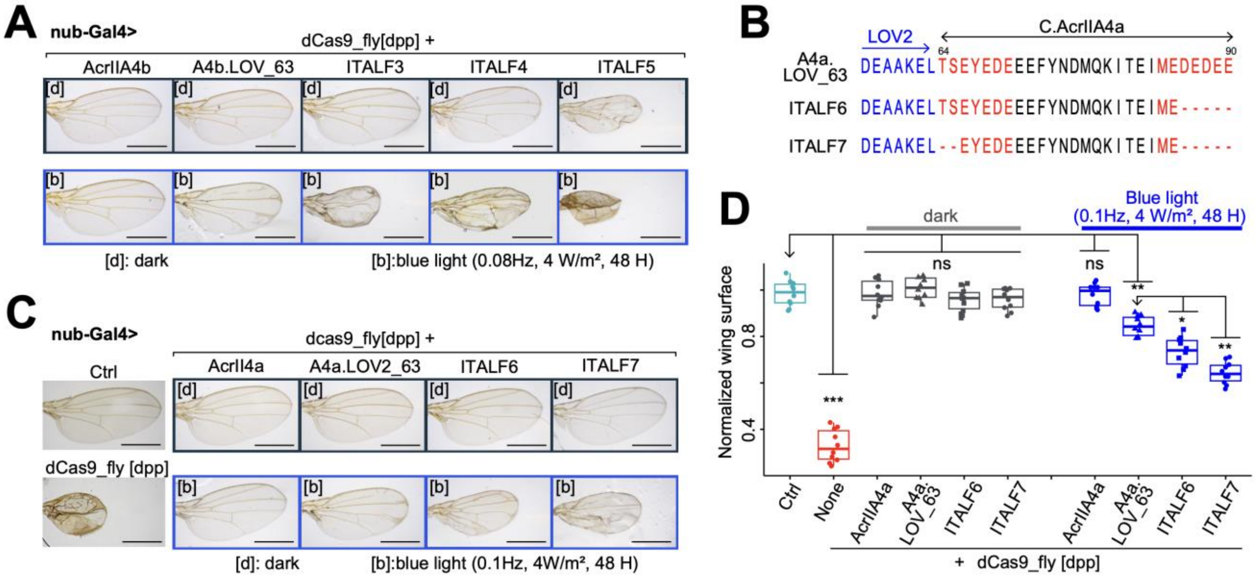


**Figure S8**: **Effect of truncation of IOR on photoswitchability of AcrIIA4.** (**A**) Representative images of wing phenotypes from dCas9_fly flies, whose larvae were kept in the dark or exposed to a blue-light, expressing Acr-LOV2 complexes in Figure 4K: A4b.LOV_63, ITALF3, ITALF4 and ITALF5. (**B**) Partial sequence graphs of ITALF6 and ITALF7 aligned with A4b.LOV_63, featuring their truncated residues ∂86-90 and ∂64/65/86-90, respectively. (**C**) Representative images of wing phenotypes highlighting the difference between dCas9_fly flies expressing full length AcrIIA4a-LOV2, ITALF6 and ITALF7. The flies were kept in the dark or exposed to blue-light pulse at the pupal stage. (**D**) Statistical data from **C**. *P* values were calculated by ANOVA-test with n = 10 females; *ns*, *, ** and *** are not significant, <0.01, <1e-3 and <1e-5, respectively, compared to that of normal wing control. In **A** and **C**, the scale bars are 500 μm.

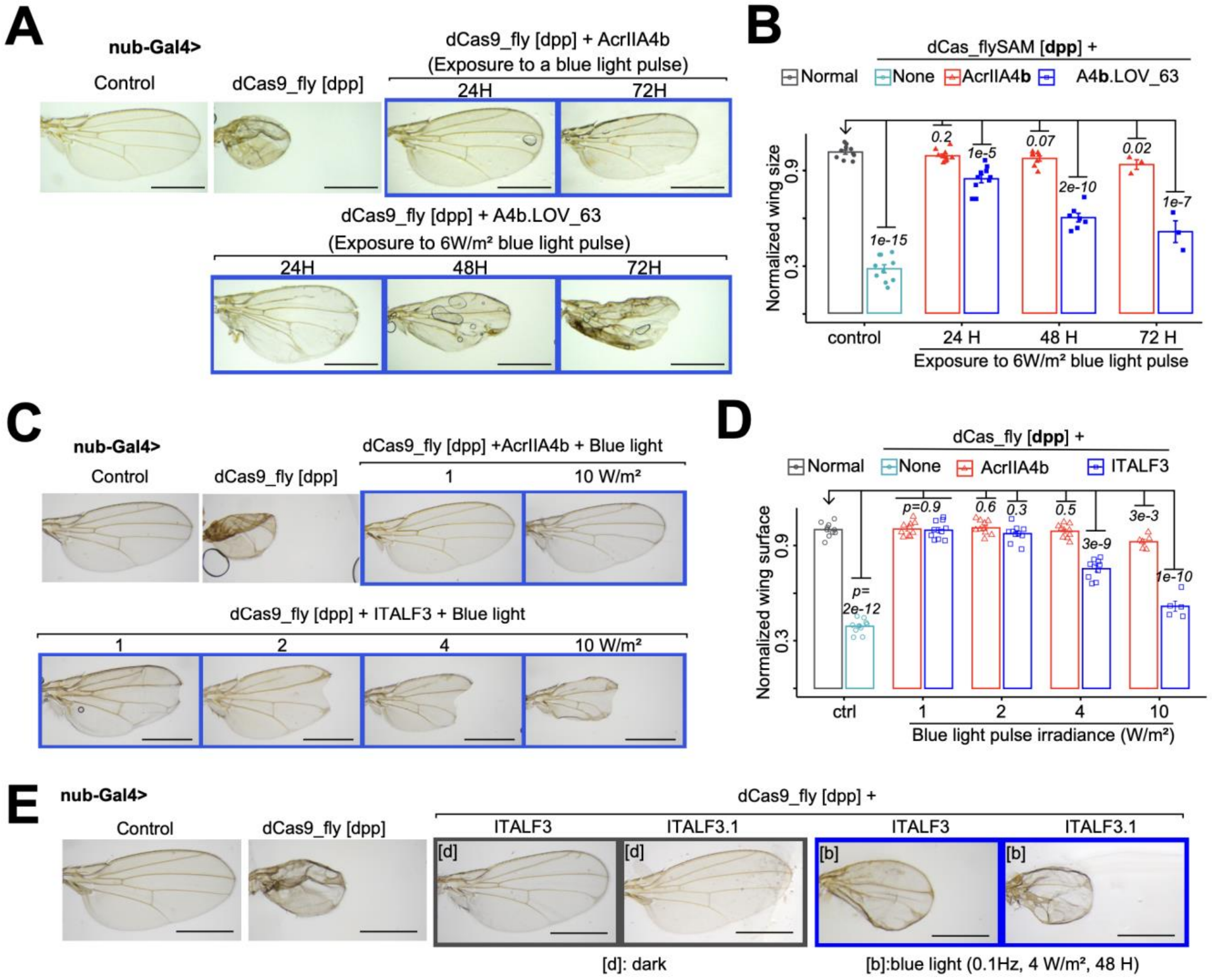


**Figure S9**: **Adjustment of light properties.** (**A-B**) Delimitation of exposure time to blue light pulse. Representative images (**A**) and statistical data (**B**) of wing phenotypes from dCas9_fly drosophila, expressing AcrIIA4b or A4b.LOV_63, exposed to light (0.2Hz, 6W/m$^2$) for different duration at a larval stage. (**C-D**) Determination of optimum light irradiance. Representative images (**C**) and statistics (**D**) of wing phenotypes from dCas9_fly drosophila, expressing AcrIIA4b or ITALF3, exposed at different irradiance of light (0.2Hz, for 48H) at a pupal stage. (**E**) Representative images of wing phenotypes from dCas9_fly drosophila, expressing ITALF3 or ITALF3.1, and the larvae of which were exposed to blue light (0.1Hz, 4W/m$^2$, 48H). In **A, C** and **E**, the scale bars are 500 µm. In **B** and **D**, the *P* values on the top of each bar were calculated by ANOVA-test of the difference between the normalized wing sizes of control flies and every group.

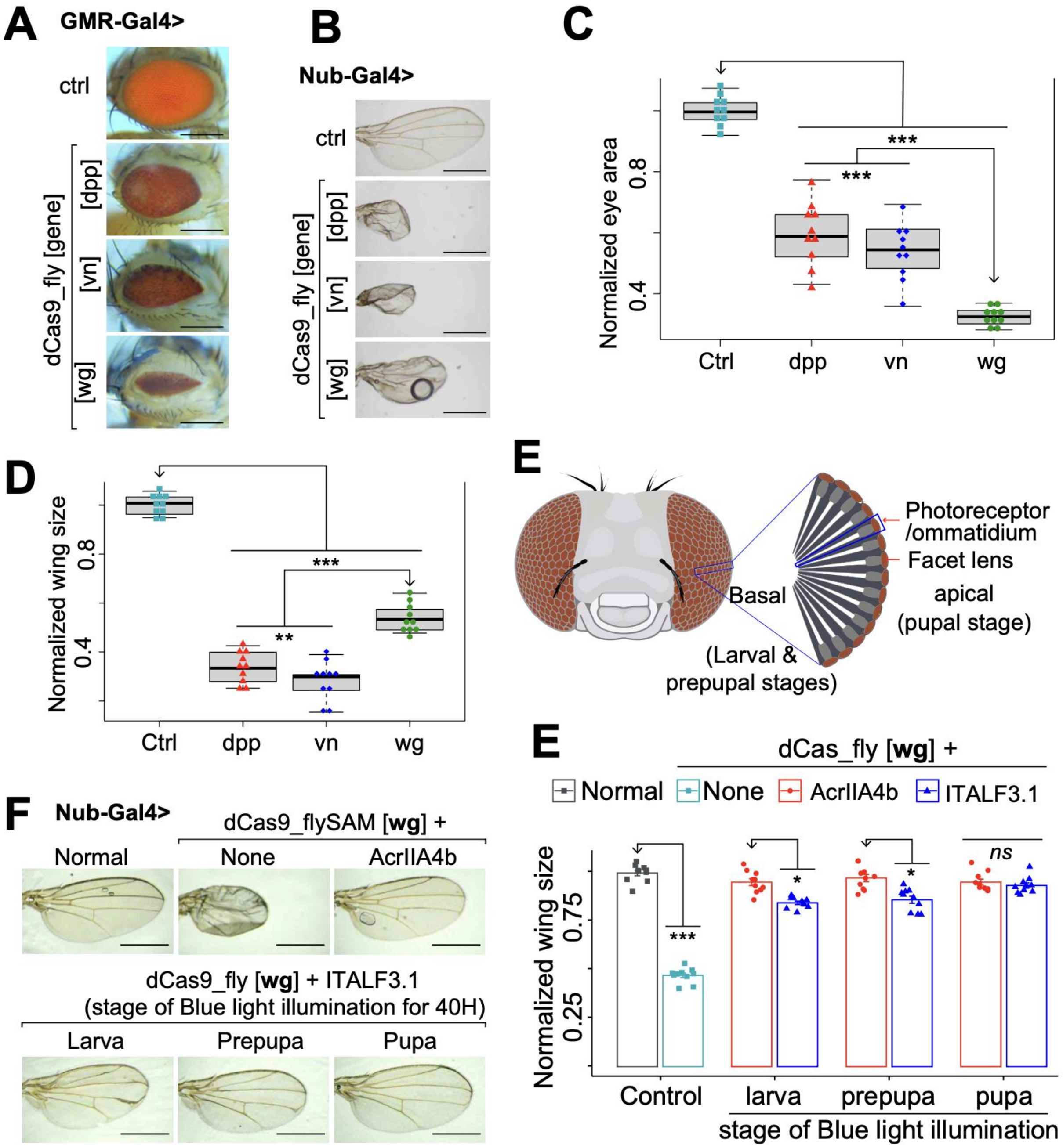


**Figure S10**: **Phenotypic changes induced by the activation of *dpp*, *vn* and *wg* genes.** (**A-B**) Representative images of GMR-Gal4 (A) and Nub-Gal4 (B) driven overexpression of *dpp*, *vn* and *wg* in eyes and wings, respectively. (**C-D**) Boxplots represent the statistical data from A in (**C**) and B in (**D**). They display that overexpression of *wg* is ideal for eye modelling, while *dpp* and *vn* for wing. (**E**) Simplified morphology of drosophila photoreceptors and the effects of *wg* overexpression on their development. (**F-G**) Representative images of wing phenotypes (**F**) and visualized statistics (**G**) from dCas9_fly flies, expressing ITALF3.1 or AcrIIA4b and. The dCas9_fly activated *wg* gene; the larvae, prepupa or pupae of these flies were irradiated with blue light (0.1Hz, 4W/$m^2$) for 30H. *P* values calculated by ANOVA-test are the mean of normalized wing sizes from dCas9_fly flies expressing ITALF3.1 compared to those expressing AcrIIA4b. The scale bars are 200 µm in **A** and 500 µm in **B** along with **E**. In **C, D** and **F**, the *p* values * < 0.01, ** <1e-3, *** < 1e-5 were calculated by one-way analysis of variance (ANOVA), with n = 10 female flies, between the normalized phenotypes of the arrowhead-pointed group and those of other groups.
